# Dynamic reorganization and selective segregation of mitochondria under DarT-mediated mtDNA damage

**DOI:** 10.1101/2022.02.08.479562

**Authors:** Nitish Dua, Akshaya Seshadri, Anjana Badrinarayanan

## Abstract

Mitochondria are dynamic organelles that play essential roles in cell growth and survival. Processes of fission and fusion are critical for distribution, segregation and maintenance of mitochondria and their genomes (mtDNA). While recent work has revealed the significance of mitochondrial organization for mtDNA maintenance, impact of mtDNA perturbations on mitochondrial dynamics remains less understood. Here we develop a tool to induce mitochondria-specific DNA damage, using a mitochondrial-targeted base modifying bacterial toxin, DarT. Following damage, we observe dynamic reorganization of mitochondrial networks, likely driven by mitochondrial dysfunction. Changes in organization are associated with loss of mtDNA, independent of mitophagy. Unexpectedly, perturbation to exonuclease function of mtDNA replicative polymerase, Mip1, results in rapid loss of mtDNA. Our data suggest that, under damage, partitioning of defective mtDNA and organelle are de-coupled, with emphasis on mitochondrial segregation independent of its DNA. Together, our works underscores the importance of genome maintenance on mitochondrial function, that can act as a modulator of organelle organization and segregation.

## Introduction

Mitochondrial networks are highly dynamic and undergo changes in shape and size throughout the cell cycle (Westermann, 2010). These dynamics are predominantly attributed to two processes of fission and fusion, carried out by evolutionarily conserved machineries (Westermann, 2008). In budding yeast, fission of mitochondria is carried out by Dnm1, which is recruited to the outer membrane by adaptor proteins. Fusion of mitochondria occurs at the level of the inner membrane (carried out by Mgm1) as well as the outer membrane, via Fzo1 (Scott & Youle, 2010; Shaw & Nunnari, 2002). Fission and fusion are required for proper distribution and segregation of mitochondria during cell division and also essential for moulding the organelle network in response to changing growth conditions (Detmer & Chan, 2007). Along with facilitating dynamic adaptation of mitochondrial function to the metabolic needs of the cell (Bagamery et al., 2020; Laporte et al., 2018), they play a crucial role in quality control of the organelle (Detmer & Chan, 2007; Ni et al., 2015).

Apart from ensuring organelle health, fission and fusion are also important for maintaining mitochondrial genome integrity and its DNA copy numbers (Busch et al., 2014; Lewis et al., 2016; Lieber et al., 2019; Osman et al., 2015). All eukaryotic cells, with a few exceptions, contain multiple copies of mtDNA (chiefly encoding for part of the electron transport chain as well as mitochondrial translation RNAs and proteins) and are distributed throughout the organelle network (Brown et al., 2011; Chen & Butow, 2005; Osman et al., 2015). These DNA copies are packaged in the form of nucleoids (Farge et al., 2012), primarily by Abf2 in budding yeast (Chen & Butow, 2005). Studies have shown that mtDNA nucleoids are non-randomly distributed in the mitochondrial network, and are present equidistant from each other (Osman et al., 2015). This organization appears to be essential for ensuring segregation of accurate amounts of mtDNA during cell division (Jajoo et al., 2016). Indeed, lowered levels of mtDNA replication are observed when mitochondrial fusion is perturbed, leading to a decrease in mtDNA copy numbers (Ramos et al., 2019). Mitochondrial fission is also linked to mtDNA replication and segregation in budding yeast and facilitates distribution of mtDNA throughout the mitochondrial network (Lewis et al., 2016). Previous studies have also reported that deletions on mtDNA occur when pathways of fission and fusion are perturbed (Osman et al., 2015).

Accurate maintenance of mtDNA integrity is critical, as mutations on mtDNA can have adverse effects on cell survival and are associated with several human pathologies (Hahn & Zuryn, 2019; Scheibye-Knudsen et al., 2015; Taylor & Turnbull, 2005). Close proximity of mtDNA to the electron transport chain makes it susceptible to damage via reactive species generated during oxidative phosphorylation (Alexeyev et al., 2013; Yakes & Van Houten, 1997). Even under physiological conditions, mtDNA accumulates mutations and deletions over time; these are also proposed to be major drivers of ageing (Alexeyev et al., 2013; Richter, 1995; Wallace, 2010).

How do cells respond to mtDNA damage? There is evidence for the localization of certain DNA repair proteins to the mitochondria (such as those involved in base excision repair and recombination) (Alexeyev et al., 2013; Boesch et al., 2011; Scheibye-Knudsen et al., 2015; Stevnsner et al., 2002). However, certain repair pathways seem to be absent from the mitochondria (for example, nucleotide excision repair) (Clayton et al., 1974; Pascucci et al., 1997). Thus, while DNA repair pathways have been detected in the mitochondria, importance and regulation of mtDNA damage repair has remained unclear. Since mtDNA can directly affect functionality of the organelle, it is possible that one layer of quality control is carried out by mechanisms that respond to mitochondrial dysfunction directly. Indeed, in response to organelle dysfunction, cells employ fission and fusion machineries to clear out any damaged mitochondria (Ni et al., 2015). For example, regulation of fusion can facilitate complementation of mutations present on mitochondrial genomes (Nakada et al., 2001). Similarly, mitochondrial fission helps in selective removal and degradation of mitochondria via PINK-Parkin mediated mitophagy or even mitocytosis (Jiao et al., 2021; Jin & Youle, 2012; Zhang et al., 2019). More recently, a role for mitochondrial reorganization at the level of the cristae has also been discovered to contribute to segregation of defective mitochondrial domains, arising from mutant mtDNA copies (Jakubke et al., 2021). It is likely that these pathways contribute to different degrees of quality control, depending on the cell type and specific growth environments (Pickles et al., 2018). Although it is becoming increasingly evident that mitochondrial dynamics can regulate mtDNA synthesis, partitioning and homeostasis, the influence of mtDNA damage on mitochondrial organization and segregation is not fully understood. A major challenge in studying mtDNA damage response and repair has been the ability to generate mtDNA-specific damage. In this context, tools to make mtDNA-specific DSBs have been invaluable, but are dependent on the presence of specific sequences on the mtDNA to allow for targeted action of the enzyme. In most cases, generic damaging agents that can cause adducts or lesions on DNA (nuclear and mitochondrial) preclude the ability to disentangle mitochondria-specific effects.

To characterize the impact of mtDNA perturbations on organelle organization and function, we developed a novel system to generate inducible mtDNA-specific damage in *S. cerevisiae*. We engineered a mitochondrially-targeted bacterial toxin, mtDarT (which generates adducts on single-stranded DNA) (Jankevicius et al., 2016) to probe the effect of mtDNA damage on mitochondrial dynamics in real-time. We find that mtDNA damage drives network fragmentation; based on our observations, we suggest that this is likely due to a lowering in fusion rates arising from mitochondrial dysfunction. We further implicate a critical role for the exonuclease function of the mtDNA polymerase in mtDNA maintenance under damage. Our study reveals an intricate connection between mtDNA integrity, mitochondrial function and its role in regulating organelle organization and segregation.

## Results

### System to induce mitochondria-specific DNA damage

To perturb mtDNA integrity and observe the resultant mitochondrial dynamics in real-time, we developed an mtDNA-specific damage inducing system in *S. cerevisiae*. For this, we utilized a bacterial toxin, DarT, that is part of a widely conserved DarT-DarG toxin-antitoxin system (Jankevicius et al., 2016). DarT functions via transferring an ADP-ribosyl group on single-stranded DNA using NAD+ as a substrate (Fig. 1A) and has been shown to generate DNA damage-inducing adducts on the bacterial genome (likely via blocking replication progression) (Lawarée et al., 2020). The adduct is attached to a thymidine within a ‘TNTC’ motif (Jankevicius et al., 2016; Schuller et al., 2021)., which is present abundantly on the mitochondrial genome.

**Figure 1:**
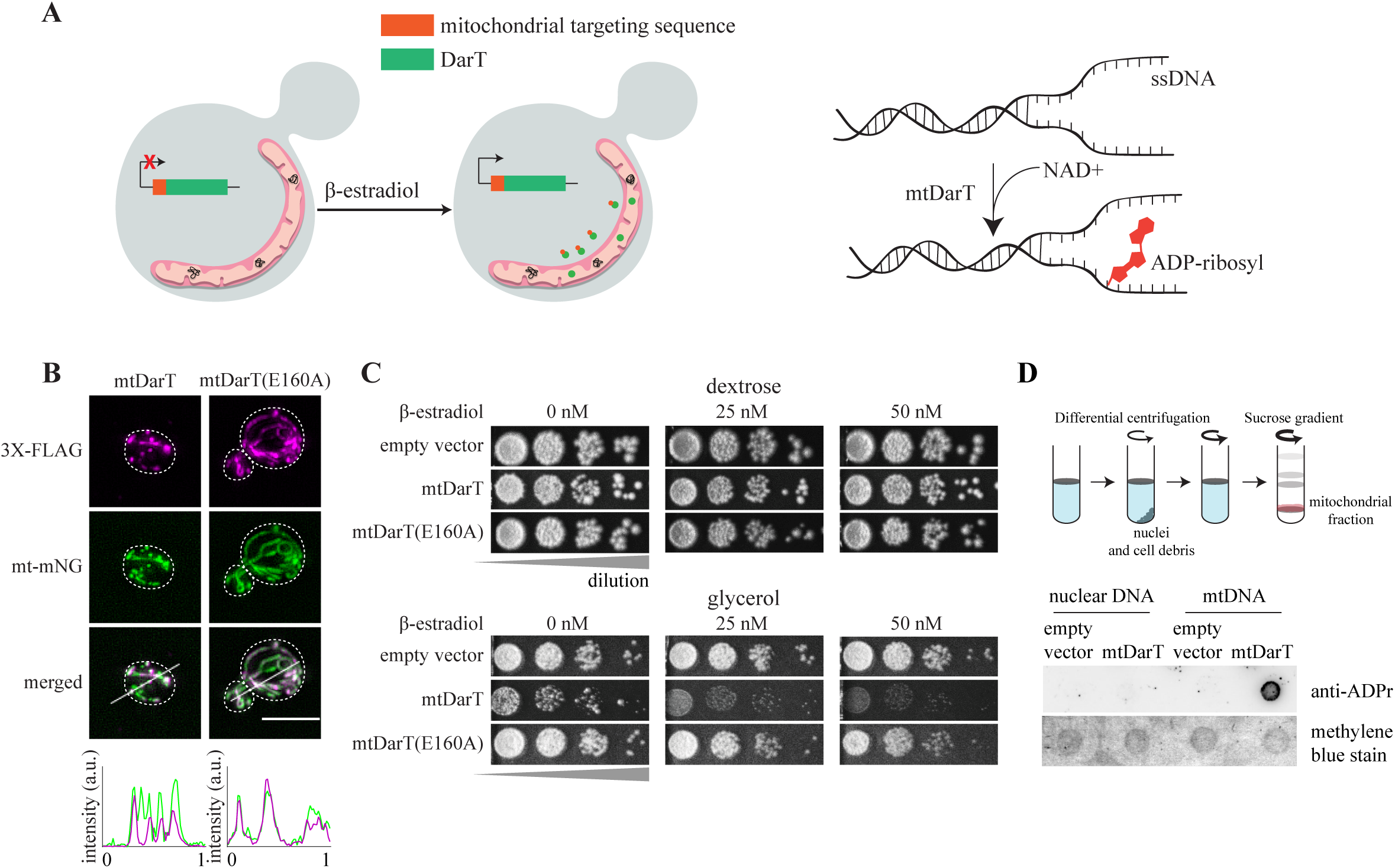
System to induce mitochondria-specific DNA damage. (A). Schematic representing expression and localization of mtDarT on addition of β-estradiol (Left panel); Diagrammatic representation of ADP-ribosylation of single-stranded DNA in presence of DarT and NAD+ (Right panel). (B). Immunostaining for FLAG-tagged mtDarT/TE160A in cells expressing mitochondrially-targeted mNeongreen. Intensity line profiles for mtDarT and mNeonGreen are show at the bottom. (Scale bar-5μm, white dotted lines mark cell outlines, here and in all other images) (C). Cell survival under increasing concentrations of β-estradiol on plates containing either dextrose or glycerol as a carbon source; cells were transformed with plasmids expressing mtDarT/TE160A or empty vector as a control. (D). Dot blot assay to assess ADP ribosylation on DNA, upon mtDarT expression. Both mitochondrial and nuclear DNA is probed. As a loading control, same volumes of samples were spotted on a separate membrane and stained with 0.04% of methylene blue

We constructed a mitochondrial matrix-targeted DarT (derived from *T. aquaticus*), mtDarT, via fusion of DarT to the first 69 amino acids of SU9 (subunit 9 of the F_1_-F_0_ ATPase) (Horie et al., 2003). A 3XFlag-tag was added to the C-terminus of DarT, to enable verification of expression as well as localization upon induction (Fig. S1A). mtDarT was expressed from a β-estradiol inducible-promoter on a low copy plasmid (Louvion et al., 1993). This strategy was employed to separately express a catalytically dead variant of DarT (DarTE160A), which retains only negligible DarT activity (Jankevicius et al., 2016). We first confirmed expression of mtDarT, as well as mtDarTE160A via western blotting after 4 and 8hr of induction with 100nM β-estradiol (Fig. S1B). Immunostaining using an anti-flag antibody revealed that the proteins indeed localized to the mitochondria, marked using mitochondrially-targeted mNeonGreen (mt-mNG) (Fig. 1B, S1C).

We next assessed the impact of mtDarT on cell growth, utilizing the ability of yeast cells to show differential growth on fermentable/ non-fermentable carbon sources, based on their mitochondrial activity (Ephrussi et al., 1955). Cells with impaired mitochondrial function are able to survive on media containing a fermentable carbon source (albeit forming smaller, ‘petite’ colonies), due the ability of these cells to carry out glycolysis. However, such cells are compromised for growth on non-fermentable carbon sources, where oxidative phosphorylation (and hence complete mitochondrial function) is essential. When compared with an empty vector control, we found that cells expressing mtDarT showed no associated loss in viability on plates supplemented with dextrose (fermentable carbon source) (Fig. 1C). These cells showed significant loss of viability in case of plates supplemented with glycerol (non-fermentable carbon source) (Fig. 1C). Importantly, the growth defects observed were mtDarT-specific and required the DNA adduct-forming catalytic function of DarT (Fig. 1C).

To further establish mitochondria-specific mtDarT effects, we carried out a dot-blot assay, to probe for ADPr groups on both nuclear and mitochondrial DNA. We observed signal only for mtDNA, dependent on mtDarT expression (Fig.1D). In addition, we assessed the localization of Lcd1p (or Ddc2p), that is reported to localize to the nucleus under nuclear DNA damage or replication stress ((Lisby et al., 2004; Melo et al., 2001); Fig. S1D, HU-treated cells). Upon mtDarT induction, we did not observe an increase in the number of cells with Lcd1p localizations (Fig. S1D). Together, these results supported the idea that mtDarT had mitochondria-specific action.

### mtDNA damage impacts mitochondrial organization and dynamics

We next characterized the impact of mtDNA damage on mitochondrial organization and function in glycolysis replete conditions. For this, we followed mitochondrial dynamics in cells expressing Cox4-mCherry to track mitochondria in addition to mtDarT (or an empty vector control). We classified the observed mitochondrial organizations into three distinct categories: ‘tubular’, ‘fragmented’ and ‘aggregate’ (representative examples are shown in Fig. 2A). We found that the induction of mtDarT resulted in fragmentation of the mitochondrial network (Fig. 2A-B), with an increase in percentage of cells with fragmented or aggregate mitochondria. Indeed, cells carrying mtDarT displayed some perturbation to mitochondrial organization even prior to induction, likely due to leaky expression from the vector (Fig. S2A).

**Figure 2:**
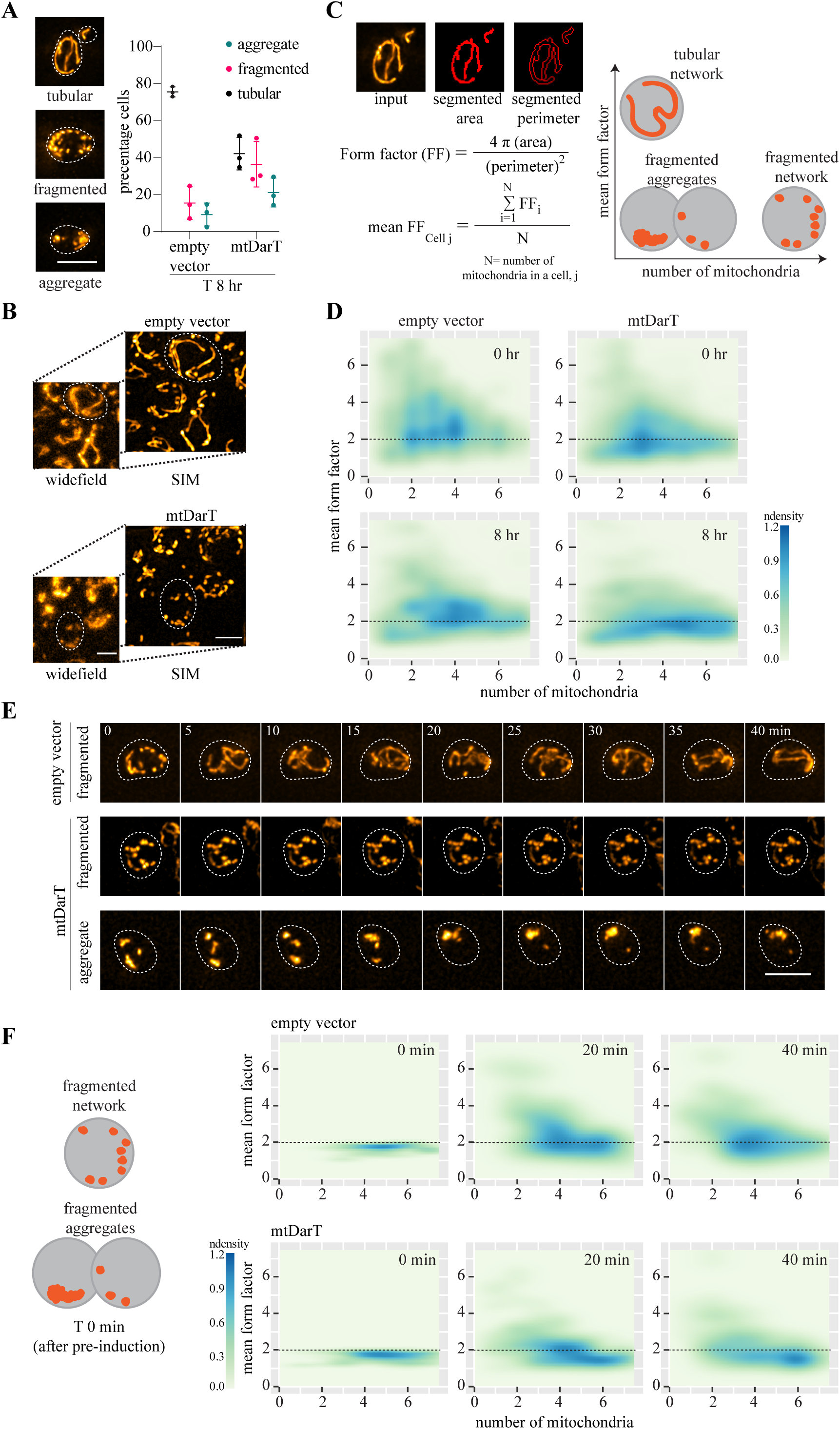
mtDNA damage impacts mitochondrial organization and dynamics. (A). Representative cells showing three categories of mitochondrial morphologies observed (left panel); Percentage of cells in each category in control (empty vector) or mtDarT cells after 8 hr of induction (Mean ± SD from three biological replicates, n>150 in each replicate) (right panel). (B). Cells expressing Cox4::mCherry imaged in widefield and SIM mode, with representative cells showing tubular and fragmented mitochondrial morphologies after 6 hr of induction with 100nM β-estradiol. (C). Two parameters: - Mean form factor (MFF) and number of mitochondria are used to quantify mitochondrial morphologies in the population. Details on calculation of MFF is provided, along with a diagrammatic representation of possible mitochondrial organizations observed in the 2D density plots (D). Density plots with Mean form factor and number of mitochondria over time; n>680 in each plot, from 3 biological replicates. (E). Montage of cells with Cox4::mCherry, imaged every 5 min for 40 min, after 4 hr of induction with β-estradiol; mitochondrial morphology at the start of the experiment is indicated. T0 refers to the frame at the start of the imaging (after the period of pre-induction) here and in Fig. 2F. (F). Density plots with Mean form factor and number of mitochondria over time, plotted for cells starting with MFF<2 at the beginning of time lapse; n=84 for empty vector, n= 151 for mtDarT pooled from 3 biological replicates.

We next followed the mtDarT-induced changes in mitochondrial organization over time via measuring two features (Fig. 2C): a. the mean form factor (MFF) of mitochondria per cell, an indicator of mitochondrial circularity (Chaudhry et al., 2020) and b. the number of discrete mitochondria per cell. We plotted MFF and number of mitochondria for all cells as 2D density plots, allowing us to visualize populations of cells having either tubular, fragmented or aggregated mitochondria (Fig. 2C). We observed that control cells harboring an empty vector largely populated MFF above 2 (predominantly tubular mitochondrial morphology) and maximum density represented an intermediate between a tubular network and fragmented state (Fig. 2D). This is as expected under unperturbed conditions, given the dynamic nature of the mitochondria throughout the cell cycle (Shaw & Nunnari, 2002). In contrast, upon mtDarT induction, mitochondria gradually transitioned from tubular (MFF >2) to fragmented/ aggregate states, having MFF <2 (Fig. 2D).

Organization of the mitochondrial network is regulated by a dedicated set of GTPases that control mitochondrial membrane fission and fusion (Westermann, 2008). Hence, to assess the contribution of changes in mitochondrial fission and/ or fusion upon mtDNA damage, we followed mitochondrial dynamics via time-lapse microscopy. We imaged cells at 5 min intervals for 40 min, after 4hr of damage exposure (Fig. 2E, S2B-C) and plotted density plots as described above. Transition from tubular to fragmented states would reveal fission events, while transition in the opposite direction (towards a tubular network) would be indicative of fusion events.

To assess the impact of mtDNA damage on fission dynamics, we followed cells (control and damage) with tubular mitochondrial networks at the start of imaging (T0), after pre-induction. We found that the rates of mitochondrial fission (as assessed by transition of cells with MFF >2 to MFF <2), were comparable between control and damage-treated cells (Fig. S2D). In line with this observation, we did not find an increase in levels of the fission protein Dnm1 (Bleazard et al., 1999) (Fig. S2E). Another mechanism for mitochondrial fission, independent of changes in the levels of Dnm1, is via ERMES contact sites (Lewis et al., 2016). An increase in the number of these contact sites (which can be imaged via tracking the number of Mdm34 localizations) could lead to recruitment of Dnm1, thereby resulting in an increase in fission events. However, we did not observe a significant increase in the number of ERMES contact sites in cells facing damage (Fig. S2F).

Unlike the observed fission dynamics, cells facing mtDarT-induced damage had significantly lower fusion events. When we followed individual cells over time, we found that mtDarT-expressing cells with fragmented or aggregate mitochondrial organization did not display dynamic reorganization towards a network morphology (Fig. 2E, S2C). Consistently, in comparison to the control, cells expressing mtDarT were compromised in transitioning from MFF <2 to MFF >2 (fragmented to tubular transition) and tended to maintain the density distribution observed at the start of the experiment (Fig. 2F). The control cells on the other hand were able to dynamically transition from MFF <2 to MFF>2 within the 40 min imaging interval (Fig.2E-F). Importantly, the above described effects of mtDarT were not cell size or cell cycle-stage specific (as assessed by the presence and size of the bud), further underscoring the direct impact of mtDNA damage on mitochondrial organization (Fig. S3A-B).Taken together, our data suggest that mtDNA damage drives a large scale reorganization of the mitochondrial network, likely via an impact on mitochondrial fusion rates. Indeed, although we did not observe a significant change in the levels of fusion protein, Fzo1p (Fig. S2E, S2G), it is possible that other mechanisms contribute to the observed mitochondrial dynamics under damage (see Discussion).

### mtDNA damage results in mitochondrial dysfunction

Given our observations on the impact of mtDNA damage on mitochondrial membrane fusion dynamics, we wondered whether mitochondrial function was perturbed in these cells. We thus measured two indicators of mitochondrial activity: a. mitochondrial membrane potential (ψ) via imaging TMRE (a membrane potential sensitive dye) and b. oxygen consumption rate (OCR) using a Seahorse assay (Ludovico et al., 2001). We observed a decrease in membrane potential in cells expressing mtDarT (Fig. 3A), similar to impact of FCCP, a membrane decoupler which decreases ψ (Fig. S4A). Similarly, mtDarT-treated cells also had lower OCR (Fig. S4B). Together, these observations suggest that mitochondrial function in mtDarT-treated cells is compromised (we use the term ‘dysfunction’ to encompass both these parameters of mitochondrial function). Importantly, lowering in membrane potential was observed even in mtDarT cells with reticular/ tubular networks, suggesting that mitochondrial dysfunction upon mtDNA damage preceded changes in mitochondrial organization and dynamics (Fig. S4D).

**Figure 3:**
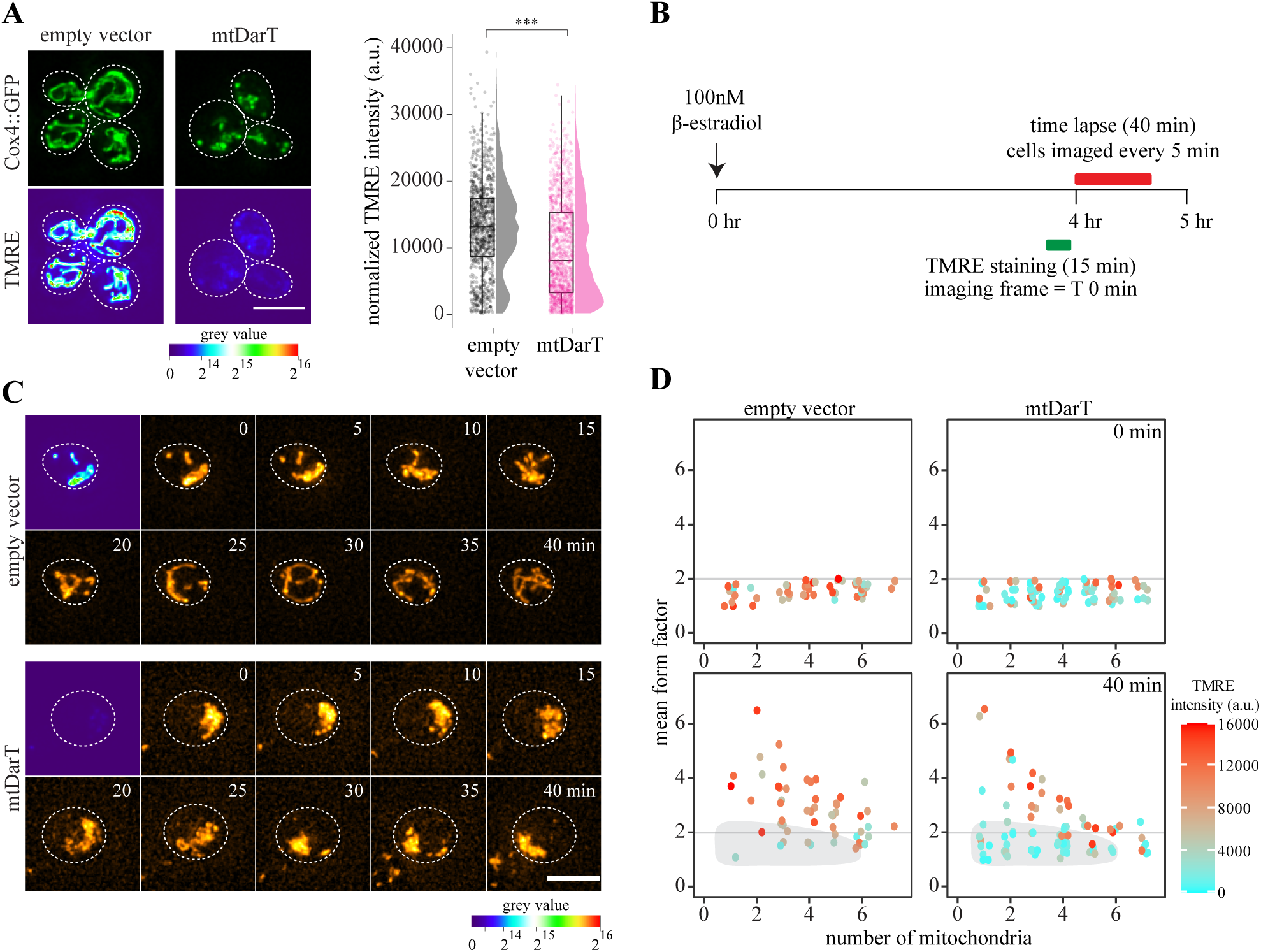
mtDNA damage results in mitochondrial dysfunction. (A). Representative images of cells expressing Cox4::GFP and stained with TMRE after 8 hr of induction with β-estradiol (left panel). Total TMRE intensity normalized to cell area is plotted (n>1030 from 3 biological replicates) (right panel). (B). Schematic of experimental design for time lapse experiment in Fig. 3C-D. (C) Montage of cells expressing mt-mNeongreen stained with TMRE tracked over time, imaged every 5 min for 40 min. TMRE was imaged only at t=0 of time lapse. T0 refers to the frame at the start of the imaging (after the period of pre-induction). (D) Scatter plot with Mean form factor and number of mitochondria over time, plotted for cells starting with MFF<2 at the beginning of time lapse. Each data point represents one cell with colour corresponding to TMRE intensity normalized to cell area at t=0 min; n=61 for empty vector, n=91 for mtDarT from 3 biological replicates. Statistical significance in A was calculated using Mann-Whitney U test. (*** p<0.0001, ** p<0.005, * p< 0.05, n.s.- not significant, in all cases).

Specific lowering in mitochondrial fusion rates has been reported in cells with compromised membrane potential (and hence dysfunction) (Duvezin-Caubet et al., 2006; Karbowski et al., 2004; Legros et al., 2004; Twig et al., 2008). We thus hypothesized that mtDNA damage-induced mitochondria with lowered membrane potential would display lower probability of fusion, when compared with mitochondria with unperturbed membrane potential. To assess this, we tracked mitochondrial dynamics (via mt-mNeongreen) in mtDarT-treated cells after a brief pulse of TMRE staining (Fig. 3B-C). We then analysed the mean form factor and number of mitochondria for cells specifically having MFF<2 (fragmented) along with their initial TMRE fluorescence intensities, and asked whether these mitochondria were able to transition to a tubular state. We observed that mitochondria with higher membrane potential, but aggregate/ fragmented organization (MFF <2) were able to fuse into tubular mitochondrial networks (MFF >2) (Fig. 3C-D). In contrast, fragmented mitochondria with lower membrane potential were unable to do so during the course of imaging (Fig. 3C-D).

Our observations are consistent with the possibility that mtDNA damage results in mitochondrial dysfunction (reduced membrane potential and compromised OCR), which in-turn affects mitochondrial dynamics via impairing fusion.

### Changes in mitochondrial morphology accompany loss of mtDNA but not mitochondria

We next assessed the effects of mtDarT damage on mtDNA, and whether the observed mitochondrial dynamics contributed to clearance of damaged DNA/ dysfunctional mitochondria. Following a previously-described method, we stained cells with SYBR Green I to visualize mtDNA nucleoids (Jajoo et al., 2016) (Fig. 4A). In the control sample without mtDNA damage, less than 2% cells had no detectable mtDNA foci (‘ρ0’ cells) at all time points of imaging (Fig. 4B). In contrast, upon mtDarT induction, the percentage of ρ0 cells steadily increased, with ∼30% cells having no detectable foci post 8hr of damage induction (Fig. 4B). Consistent with this, we also observed an overall decrease in the number of mtDNA foci per cell upon mtDNA damage (Fig. S5A).

**Figure 4:**
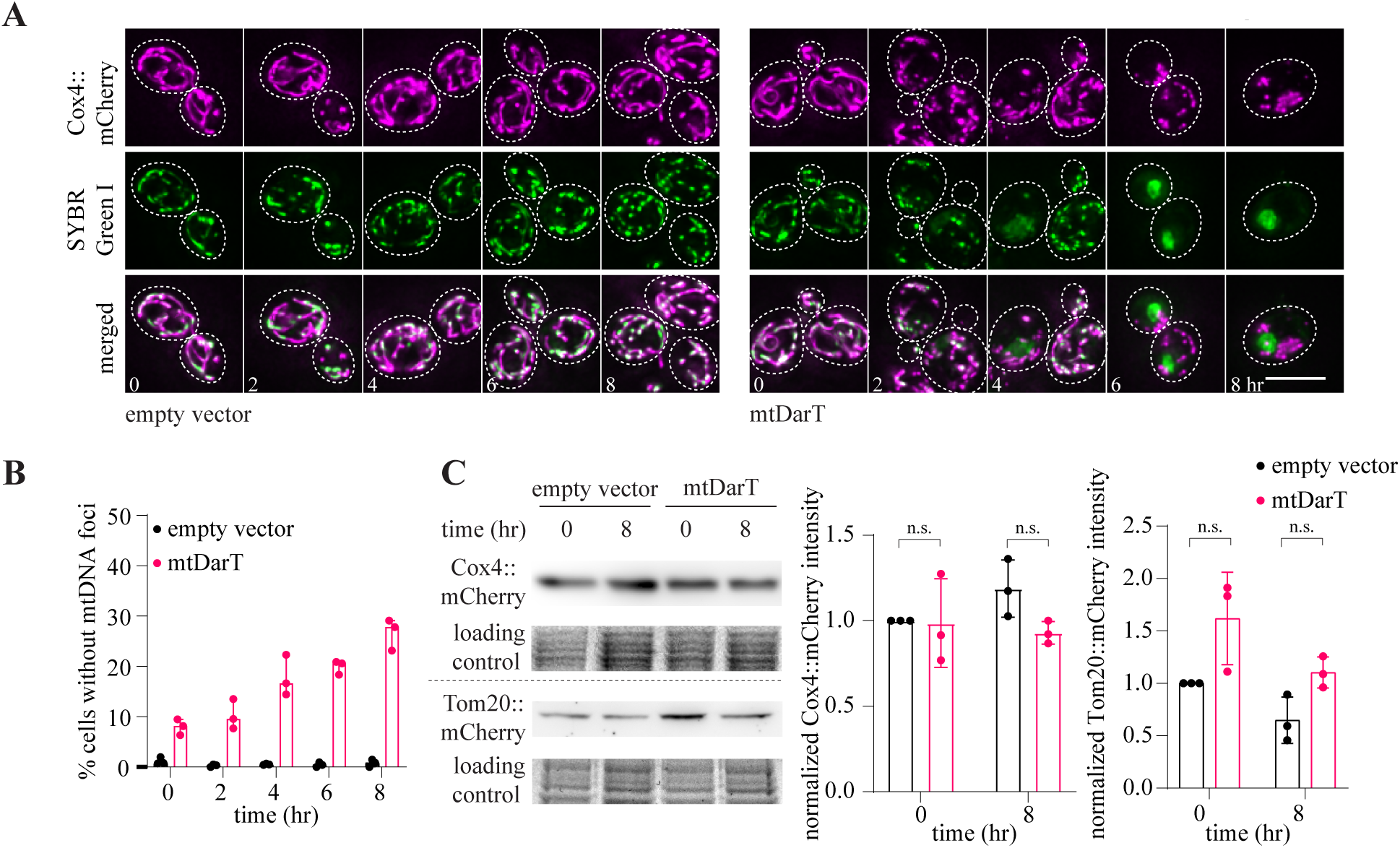
Changes in mitochondrial morphology accompany loss of mtDNA but not mitochondria. (A). Montage of cells expressing Cox4::mCherry stained with SYBR Green I to visualize mtDNA over time. (B). Percentage of cells over time without any mtDNA foci. Mean ± SD from 3 biological replicates; n>100 in each biological replicate. (C). Cox4::mCherry and Tom20::mCherry western blot with quantification from 3 biological replicates, normalized to empty vector at t=0; Error bars: SD. Statistical significance in C was calculated using unpaired Student’s t-test.

In some instance mitochondrial dysfunction (and associated fragmentation) has been shown to occur in conjunction with mitophagy (Burman et al., 2017; Kleele et al., 2021). We thus asked whether loss of mtDNA under damage was a result of clearance of dysfunctional mitochondria. For this, we assessed the levels of two nuclear-encoded mitochondrial proteins, Cox4 and Tom20, via western blot. We found that levels of these proteins did not change significantly under damage conditions (Fig. 4C). This is in contrast to mitochondrially-encoded Cox2p levels, that decreases upon mtDarT expression (Fig. S4C). We then measured mitochondrial volume (Harwig et al., 2018) (Fig. S5B) and found that here too, no significant changes were observed over time. Consistent with the lack of change in mitochondrial content, we did not observe an increase in mitophagy-associated marker, Atg8 (Torggler et al., 2017) localizations (Fig. S5C), or a significant increase in the number of Atg8 localizations associated with mitochondria (Böckler & Westermann, 2014) (Fig. S5D). We also did not detect a significant difference in the amount of vacuolar localization of Idh1 (Kanki & Klionsky, 2008) upon mtDarT-induction (Fig. S5E). This suggested that an mtDNA-specific mechanism might be responsible for mtDNA clearance in response to damage.

### Exonuclease activity of the mtDNA polymerase, Mip1, is essential for mtDNA maintenance

In mammalian cells, the mtDNA replicative polymerase PolG can switch between synthesis and degradation functions to clear out defective mtDNA (Nissanka et al., 2018; Peeva et al., 2018). Evidence for such exonuclease function of the yeast homolog, Mip1, has also been reported in conditions of starvation or oxidative stress (Medeiros et al., 2018). We hence asked whether exonuclease activity could contribute to mtDNA clearance under mtDarT-induced damage. For this, we blocked the exonuclease activity of Mip1, via mutating the catalytic site (D171A, E173A) responsible for its degradative function (Foury & Vanderstraeten, 1992; Medeiros et al., 2018) (Fig. 5A) and assessed presence of mtDNA nucleoids via SYBR Green I staining. Surprisingly, upon mtDarT induction, we observed that Mip1 exonuclease deficient cells generated ρ0 cells significantly faster than wild type. As shown earlier (Fig. 4B), 30% of wild type cells were ρ0 after 8hr of mtDarT induction. *mip1^exo-^* cells already generated ρ0 cells at a low frequency (∼7%) in no damage conditions, highlighting that inability to clear damaged mtDNA (in this case likely due to replication errors) results in mtDNA loss in glucose replete growth environment. In line with this observation, induction of mtDNA damage resulted in 60% cells becoming ρ0 post 8 hr of mtDarT induction (Fig. 5B). Importantly, rapid mtDNA loss did not result in production of cells devoid of mitochondria.

**Figure 5:**
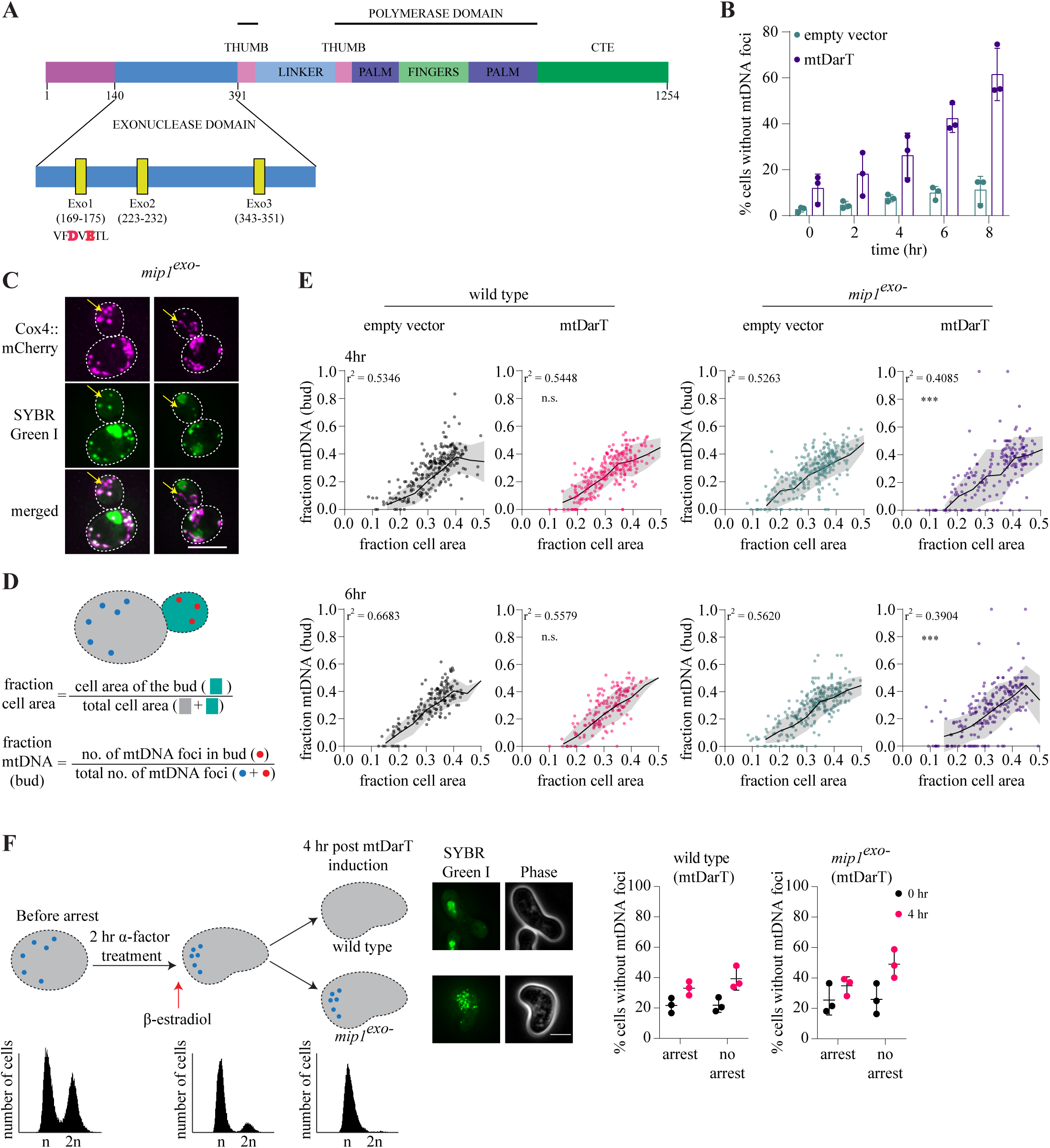
Exonuclease activity of the mtDNA polymerase, Mip1, is essential for mtDNA maintenance. (A) Schematic showing domain organization of Mip1, with amino acid mutations for *mip1^exo-^* marked in red (D171A, E173A). (B) Percentage of cells over time without any mtDNA foci (*p^0^*) are plotted. Mean ± SD from three biological replicates; n>85 in each biological replicate. (C) Representative images of *mip1^exo-^* cells expressing Cox4::mCherry stained with SYBR Green I. Yellow arrows mark mitochondria devoid of mtDNA. (D-E). Fraction of mtDNA in bud (number of mtDNA foci in bud/total number of mtDNA foci in the mother-bud pair) is plotted against fraction cell area (area of the bud/total area of the mother-bud pair) for both wild type and *mip1^exo-^* cells, after 4 hr (top panel) or 6 hr of induction (bottom panel). Black line represents mean and shaded region represents SD (calculated after binning x-axis in bins of 0.05). R^2^ values were obtained via Pearson’s Correlation; n>145 from 4 biological replicates for *mip1^exo-^* and 3 biological replicates for wild type. Statistical significance in E was calculated using Fisher’s F-test, applied on distributions of residual values of all data points (distance from linear fit curve). Comparisons were made to empty vector control of corresponding time point. (F) Schematic of experimental design to arrest cells, followed by SYBR staining at 0 and 4 hr of induction. Flow cytometry profiles corresponding to the time point are shown at the bottom (left); percentage cells without mtDNA foci in wild type and *mip1^exo-^* cells expressing mtDarT (right). N>670 cells from 3 biological replicates.

How do *mip1^exo-^* cells lose mtDNA in such a rapid manner under damage? We measured budding events of wild type and mutant cells upon mtDNA damage and found them to be comparable during the course of the experiment (3±1 budding events over 6 hr, for both wild type and *mip1^exo-^* cells expressing mtDarT), ruling out the possibility of faster division rates in the exonuclease mutant contributing to mtDNA loss over time. We next assessed mitochondrial dynamics in the mutant. As seen in wild type cells, we found that mtDNA damage resulted in significant changes to mitochondrial organization in case of exonuclease deficient cells as well (Fig. S6A). Furthermore, in comparison to mtDarT-treated wild type cells, we found that membrane potential was severely compromised in the mutant (Fig. S6B), suggesting that these cells likely have significantly lowered mitochondrial membrane fusion events. A potential outcome of enhanced mitochondrial dysfunction is a defect in the segregation of mitochondria during division (McFaline-Figueroa et al., 2011). We thus wondered whether this could impact partitioning of the mtDarT-induced damaged mtDNA. Such partitioning defect would likely be more severe/ pronounced in the *mip1^exo-^* condition, where defective mtDNA would accumulate due to lack of a clearance mechanism.

When we probed for mtDNA in these cells, we observed a significant proportion of cells with mitochondria devoid of mtDNA in *mip1^exo-^* background (Fig 5C). We asked whether mtDNA was differentially partitioned in these cells, resulting in generation of mitochondria devoid of DNA as well. For this, we analysed mother-bud pairs after SYBR Green I staining, and quantified the fraction of mtDNA foci in the bud (Fig. 5D). Consistent with previous reports (Jajoo et al., 2016; Osman et al., 2015), we found that the amount of mtDNA partitioned in the bud scaled with increasing bud area in control as well as wild type cells after mtDarT induction (Fig. 5E). However, unlike wild type cells, there was significant heterogeneity in the amount of mtDNA in the bud (across cell areas) in case of exonuclease deficient cells treated with mtDarT, suggesting imprecise mtDNA partitioning (Fig. 5E), with several instances of buds receiving no mtDNA (ρ0 daughter cells) (Fig. 5E). We hence hypothesized that if such asymmetry indeed contributed to the rapid production of ρ0 cells in the exonuclease deficient mutant, cells arrested for cell cycle progression would no longer generate ρ0 cells at the high frequency observed in case of steady state growth conditions. To assess this, we used *bar1*Δ strains treated with α-factor, that should arrest cells in G1-S transition (Breeden, 1997). We confirmed the same using flow cytometry, as well as via assessing the morphology of the cells upon arrest (Fig. 5F (left) and (Rosebrock, 2017)). In wild type cells, we observed that loss of mtDNA occurred at comparable frequency in arrest and non-arrest conditions. In contrast, in *mip1^exo-^* background, mtDNA loss was significantly lower under arrest (Fig. 5F (right)). Thus, asymmetric mitochondrial partitioning during cell division could significantly contribute to the rapid production of ρ0 cells in the exonuclease deficient mutant. Taking these observations together, our results suggest that mitochondrial dynamics could differentially impact damaged mtDNA segregation in conditions where defective mtDNA clearance mechanisms are compromised, likely facilitating organelle segregation independent of its DNA.

## Discussion

In this study, we describe a system to generate mtDNA-specific damage in *S. cerevisiae*. Using a bacterial toxin DarT (Jankevicius et al., 2016), targeted to the mitochondria, we follow the impact of mtDNA damage on mitochondrial organization, segregation and function. We find that mtDNA damage affects mitochondrial function via compromising membrane potential. This could be the causative force in mitochondrial reorganization by reduction in mitochondrial membrane fusion events (Duvezin-Caubet et al., 2006; Karbowski et al., 2004; Legros et al., 2004; Twig et al., 2008), facilitating selective segregation of mitochondria without defective mtDNA (discussed below).

How cells sense and respond to mtDNA damage remains an outstanding and exciting question. Previous studies have detected the presence of specific DNA repair enzymes (include BER and recombination machinery) in the mitochondria (Alexeyev et al., 2013; Boesch et al., 2011; Scheibye-Knudsen et al., 2015; Stevnsner et al., 2002). While some of these pathways encode mitochondrially-targeted isoforms (Swartzlander et al., 2010), many do not carry any known localization signal (Stuart et al., 2005; Thyagarajan et al., 1996). How they localize to the mitochondria remains unclear, as is the importance of specific repair pathways in mtDNA maintenance (Alexeyev et al., 2013; Scheibye-Knudsen et al., 2015). Indeed, there are certain types of damages (such as UV-induced pyrimidine dimers) for which active repair pathways have not been found in mitochondria (Clayton et al., 1974; Pascucci et al., 1997). In such scenarios, where mtDNA damage becomes persistent, degradation of damaged mtDNA copies has been proposed to occur (Alexeyev et al., 2013; Shokolenko et al., 2013). Interestingly, this function is carried out by the exonuclease activity of the mtDNA polymerase (Mip1 or PolG) itself (Nissanka et al., 2018; Peeva et al., 2018).

Our work further underscores a central role for defective mtDNA clearance via exonuclease activity of the replicative polymerase under damage. We propose that Mip1 exonuclease function is required for mtDNA degradation under damage and inability of cells to clear such damaged mtDNA results in rapid generation of ρ0 cells in glucose replete conditions. Although in unperturbed conditions, mtDNA and mitochondrial segregation may be coordinated (Jajoo et al., 2016; Lewis et al., 2016; Osman et al., 2015), our observations support a role for selective segregation of damaged mtDNA, in a manner that can result in partitioning of mtDNA-less mitochondria in the daughter cells. Generation of cells devoid of mtDNA, without significant loss of mitochondria would suggest cells can de-couple mtDNA and mitochondrial partitioning and maintenance under conditions of damage, when complete mitochondrial function may not be essential (such as in glycolytic conditions). Indeed, while mitochondria cannot be made *de novo* (Westermann, 2014), ρ0 and ρ-cells in the environment could regain and complement lost mtDNA, albeit with different proficiencies (Ling et al., 2019; Perlman, 1976).

It is interesting to note that mtDNA damage triggers rapid changes in mitochondrial organization and segregation, both outcomes potentially stemming from mitochondrial dysfunction. It would be important to understand how mtDNA damage causes perturbations to mitochondrial function and how this in turn modulates rates of fusion. Although levels of Fzo1 did not change under mtDNA damage, it would be informative to understand what other mechanisms might regulate mitochondrial fusion to result in the observed dynamics in budding yeast. For example, mtDNA damage can perturb levels of mitochondrially-encoded proteins such as those part of the electron transport chain (Fig. S4C), which requires coordinated effort from nuclear and mitochondrial genomes (Fontanesi et al., 2006; Herrmann & Funes, 2005). This can adversely affect maintenance of mitochondrial membrane potential. Separately, studies have reported a direct impact of lowered membrane potential on differential processing of Opa1 (a mitochondrial membrane fusion protein in mammals), affecting its association with the mitochondrial membrane (Song et al., 2007).

The change in mitochondrial organization we report is consistent with observations in mammalian cells, where mitochondrial network fragmentation has been seen to occur in response to mtDNA damage (Ghosh et al., 2019). This suggests that mitochondrial reorganization may be a universal response to mtDNA perturbation, as a fundamental quality control mechanism to deal with mitochondrial dysfunction and regulate mtDNA/ mitochondrial segregation as well as clearance. Indeed in metazoans, mitochondrial fission has been proposed to facilitate selective removal and degradation of mitochondria via processes such as PINK-Parkin mediated mitophagy (Ni et al., 2015; Scheibye-Knudsen et al., 2015). Our results instead suggest that mitophagy does not play a significant role in the mtDNA damage response in yeast (in the timescales of our present study), and that quality control mechanisms likely act at the level of mtDNA and its segregation (Fig. S5D). For example, previous reports have shown that budding yeast cells segregate mitochondria asymmetrically based on the activity of the organelle (McFaline-Figueroa et al., 2011; Vevea et al., 2014). Such asymmetry could also passively ensure selective segregation of intact mtDNA copies to daughter cells. Indeed, recent studies demonstrate selective removal or clearance of mutated mtDNA via sensing of local mitochondrial activity as well as downregulation of mitochondrial fusion (Hill et al., 2014; Jakubke et al., 2021; Lieber et al., 2019). It is tempting to speculate that in organisms like budding yeast, that divide asymmetrically, the inability of dysfunctional mitochondria to fuse with the rest of the mitochondrial network, along with selective segregation during division, is the predominant mitochondrial quality control mechanism even in response to mtDNA damage.

In sum, our study underscores: a. the impact of perturbations to mtDNA integrity on mitochondrial dynamics and function; b. the importance of Mip1 exonuclease activity in mtDNA maintenance under damage and c. the ability of cells to decouple mitochondrial and mtDNA partitioning under mtDNA damage, with an emphasis on mitochondrial maintenance even in the absence of mtDNA, in glycolysis replete conditions. Indeed, the mtDarT system developed here can provide a unique approach to enable future investigations on the mechanisms that govern mtDNA-specific damage repair and recovery across diverse growth conditions.

## Materials and methods

### Yeast strains and growth conditions

A prototrophic strain CEN.PK of *Saccharomyces cerevisiae* was used as the background strain in all the experiments (van Dijken et al., 2000). Plasmids, strains used in the study are listed in Supplementary Table 1-3. Yeast cultures were grown in rich YP media (containing 2% Peptone, 1% Yeast extract, and 2% Dextrose or 3% Glycerol as the carbon source) at 30֯ C with 200 rpm shaking with appropriate concentrations of antibiotics, as required. For all time-course and time-lapse experiments, cells were transformed with the plasmids, and transformants were inoculated in YPDextrose containing NAT and allowed to grow overnight. These cultures were induced with β-estradiol at an O.D.∼ 0.2. For induction, β-estradiol (working stock-50μM, prepared from a stock of 1mM in 100% molecular biology grade ethanol) was added to a final concentration of 100nM, unless mentioned otherwise. We performed our experiments in haploid budding yeast cells, and β-estradiol was maintained in the growth medium throughout.

Yeast transformations were done using the LiAc method. For plasmid transformations, cells were plated on YPDextrose plates containing appropriate concentrations of NAT and were allowed to grow for 48 hr at 30֯ C. *mip1^exo-^* cells transformed with plasmids, were plated on YPGlycerol plates containing appropriate concentrations of NAT and were allowed to grow for 3 days at 30֯ C.

*mip1^exo-^* mutant was generated as described earlier (Medeiros et al., 2018). Briefly, we first generated a *mip1Δ* strain which resulted in formation of 𝛒^0^ cells. We then introduced a copy of the *mip1* gene containing the mutations D171A and E173A along with a hygromycin selection cassette, at the same locus. Successful transformants containing the mutated copy of the gene were then crossed with the wild type strain of the opposite mating type to reintroduce mtDNA. The diploids were sporulated and tetrads were dissected to obtain 𝛒^+^ *mip1^exo-^* cells which were selected using hygromycin and the mating type was confirmed using PCR primers specific for mating type locus. *mip1* gene was sequenced to confirm the presence of the mutations. The strain was maintained on YPGlycerol post dissection to ensure selection of only 𝛒^+^ cells; experiments were carried out in YPDextrose.

### Dot blot assay for ADP ribosylation

For separation of mitochondrial and nuclear DNA, differential centrifugation followed by sucrose gradient based separation was employed to fractionate both the organelles. Mitochondrial fractionation was done as described earlier in (Gregg et al., 2009; Meisinger et al., 2006). DNA was isolated from the nuclear and mitochondrial fractions using a Qiagen DNeasy Blood and Tissue kit.

Isolated DNA was then used to perform a dot-blot assay as described in (Jia et al., 2017; Luviano et al., 2018). Briefly, 180ng of each DNA sample was denatured in 0.3M NaOH for 10 min at 95 ֯C. Immediately post denaturation, 5μl (∼90ng) of DNA was spotted on nitro cellulose membrane. DNA was then crosslinked to the membrane using a UV-crosslinker at 1200J. The membrane was blocked using 5% blocking solution prepared in TBST for 1 hr at RT, followed by incubation with 1:500 dilution of anti-pan-ADPr-binding reagent (Merck #MABE1016) in TBST for 1 hr at RT. The membrane was then washed thrice with TBST for 10 min each and then probed with 1:5000 dilution of HRP conjugated secondary antibody for 1 hr at RT. After the incubation, the membrane was again washed thrice, 10 min each in TBST and then developed using SuperSignal West PICO PLUS or FEMTO PLUS chemiluminescent substrate. As a loading control, same volumes of samples were also spotted on another membrane, which after crosslinking, was stained with 0.04% of methylene blue prepared in 0.5M Sodium Acetate.

### mtDNA and mitochondrial staining

For visualizing mtDNA, a previously described protocol was used (Jajoo et al., 2016). Briefly, 1ml of culture was pelleted and washed with 1X PBS. The supernatant was discarded and cells were re-suspended in the residual buffer to give 50μl cell suspension. 0.5μl of undiluted SYBR Green I (Invitrogen) was added to the cell suspension and mixed with gentle tapping. Cells were immediately pelleted and supernatant was discarded. The pellets were washed with 1ml of 1X PBS. Cells were subsequently diluted to appropriate densities in 1X PBS before imaging.

For visualizing the mitochondrial membrane potential, cells were pelleted and re-suspended in 10ml of YPDextrose media (pre-warmed to 30֯ C) containing 1.5μM of TMRE. The cells were incubated in the dark for 20 min in a shaker incubator (30֯ C, 200 rpm). Subsequently, cells were pelleted down and washed with YPDextrose media in 2X the original volume of culture used for staining. The cell pellets were resuspended to appropriate density in YPDextrose prior to imaging. For time lapses, cells were stained with TMRE post 3.5 hr of induction, in presence of 100nM β-estradiol. Following this, cells were washed and spotted on YPDextrose pads for imaging.

### Fluorescence Microscopy

Overnight cultures were allowed to grow till O.D.∼0.2. At this O.D, mtDarT expression was induced with 100nM β-estradiol. For time course experiments, samples were spotted on 1% ultra-pure agarose water pads, and for time lapse experiments, samples were spotted on YPDextrose pads with 100nM β-estradiol.

Microscopy was performed on a wide-field epifluorescence microscope (Eclipse Ti-2E, Nikon) with a 60X oil immersion objective (plan apochromat objective with NA 1.41) and illumination from pE4000 light source (CoolLED). The microscope was equipped with a motorized XYZ stage and focus was maintained using an infrared-based Perfect Focusing System (Nikon). Image acquisitions were done with Hamamatsu Orca Flash 4.0 camera using NIS-elements software (version 5.1). For time-lapse imaging, samples were placed inside an OkoLab incubation chamber maintained at 30°C and imaged at specific intervals for the indicated period of time.

For SIM imaging, coverslips were sequentially washed in methanol and 100% ethanol and flame dried prior to sample preparation. Cells were fixed in 3.7% para-formaldehyde (900μl of cell suspension was added to 100μl of 37% para-formaldehyde) and then washed with 1X PBS. Cells were then spotted on 1% ultra-pure agarose water pads and imaged. Samples were imaged with Nikon 3D-SIM microscope equipped with an ANDOR iXon DU-897 EMCCD camera (512X512), 100X ApoTIRF 1.49 NA oil objective with Coherent Sapphire 561nm solid state laser as the light source. Image reconstruction was done using NIS elements software (Illumination modulation contrast – Auto, High resolution noise suppression – 1X, Out of focus blur suppression – 0.25). During acquisition, 800ms exposure for Cox4-mCherry with 20% laser power, 150 EM gain and 0.3 μm step size was used.

### Seahorse assay

Basal respiration rate was estimated by performing seahorse flux assays using Agilent Seahorse XFe24 analyzer. 1 ml of the XF Calibrant solution was aliquoted into each well of the plate and the sensor cartridge plate was hydrated overnight as per the manufacturer’s instructions. To adhere the cells to the bottom of the wells, the plate was first coated with poly-L-Lysine for 1 hour at RT. Excess poly-L-lysine was removed and the plate was dried at 30°C. Cells grown in YPDextrose media containing appropriate concentration of NAT were induced with 100nM β-estradiol at O.D.∼0.2. Cells were serially diluted 4 hr post induction and then aliquoted into the wells such that the final number of cells in each well ∼3X10^5^. The plate was centrifuged at 100g for 2 min and incubated at 30°C for 30 min to allow the cells to adhere to the bottom of the wells. Simultaneously, the sensor cartridge was loaded with 1mM of complex IV inhibitor sodium azide and equilibrated in the Seahorse XFe24 analyzer one hour prior to the start of the assay. The plate was then loaded to the Seahorse XFe24 analyzer and the basal OCR was measured over time. A minimum of two measurements were taken with intermittent 2 min mixing and waiting steps. This was followed by sodium azide injection to the sample wells, and three measurements were taken with intermittent 2 min mixing and waiting steps.

### Arrest experiment

Cells transformed with empty vector or mtDarT plasmid were grown overnight in YPG media containing appropriate concentration of NAT. Overnight cultures were allowed to reach early log phase (O.D.∼0.2-0.6), after which they were pelleted and resuspended in pre-warmed YPD media (30^0^C) containing 20ng/ml of alpha factor with NAT. The cultures were allowed to grow at 30^0^C for 2 hr at 200rpm. Simultaneously, separate cultures without alpha factor were also maintained. Following 2 hr, samples were collected for microscopy (SYBR staining) and flow cytometry. Samples were subsequently induced with 100nM β-estradiol. Similarly, samples were collected after 4 hr of induction for microscopy and flow cytometry. (Note: Cells were pelleted and resuspended in fresh media containing alpha factor, 100nM β-estradiol and NAT 3 hr post induction to ensure continuous arrest).

### Image analysis

All image analysis was performed using MATLAB and Fiji (ImageJ) (Schindelin et al., 2012). Mitochondrial analysis was done on 2D maximum projections of z-stacks and images were deconvolved using NIS-elements AR software (Richardson-Lucy algorithm with 60-iterations). Cell segmentation was performed using a MATLAB algorithm (Doncic et al., 2013). Cell outlines were extracted from .mat output files from the algorithm and then ROIs were obtained using Fiji. For segmentation of mitochondria, a custom pipeline was set up in Fiji (briefly involving a. Background subtraction (Rolling ball radius: 50 pixels). b. Sigma Filter Plus (radius =1 pixel, use pixels within = 2 sigmas, Minimum pixel fraction = 0.2). c. Enhanced Local Contrast (CLAHE) (Blocksize = 64, histogram = 256, maximum = 2). d. Tubeness filter (sigma =1.000). e. Images converted to 8-bit. f. Thresholding (10,255) (Minimum threshold was adjusted between 8-15 depending on the quality of segmentation). g. Images converted to mask, Despeckled and Remove outliers (Radius =2, threshold =50)) and the mitochondrial descriptors (Mean form factor, mtArea, Number of mitochondria on per cell basis) were obtained using Mitochondrial analyzer (Chaudhry et al., 2020). Whole cell TMRE intensity was analyzed from maximum projections using ‘Measure’ in Fiji. For Fig. 2A, cells were manually binned into three morphological categories and buds were excluded from the analysis. For detection of mtDNA foci post SYBR Green I staining, 3D maxima finder plugin (part of the 3D ImageJSuite) in ImageJ was utilized. Since cells with less/without any mtDNA also show nuclear DNA staining, we manually scored these cells post segmentation to account for detection of foci due to nuclear staining. For mtDNA analysis in mother vs bud, pairs without any mtDNA foci (in both cell types) were removed. Mitochondrial volume was analyzed using Mitograph and volume was calculated from the number of voxels detected (Rafelski et al., 2012; Viana et al., 2015). Colocalization of GFP-Atg8 with Cox4-mCherry was scored manually as described in (Böckler & Westermann, 2014). Association of Atg8 foci with mitochondria was checked in the plane of z-stack where GFP-Atg8 intensity was maximum.

### Western blotting

Transformants were inoculated in 100-120 ml of YPDextrose with appropriate concentrations of NAT. Cultures were induced with 100nM β-estradiol at an O.D ∼ 0.2. Cells were harvested and protein isolation was performed using the Trichloroacetic acid (TCA) extraction method.

Pellets were resuspended in 300μl of 10% TCA and lysed by bead beating thrice. The precipitates were resuspended in 200μl of SDS-Glycerol buffer (7.3% SDS, 29.1% glycerol and 83.3 mM Tris base) and boiled at 95֯ C for 10 min with frequent vortexing. The lysate was precipitated and the supernatant was obtained containing protein. Estimation of protein concentration was done using the BCA estimation method using GBioSciences assay kit. The assay was performed in disposable cuvettes, where 5μl was mixed with 1ml Solution A and 20μl of Solution B and incubated at 30֯ C for 30 min. The absorbance was recorded at 565nm. Equal amounts of protein were then loaded on a polyacrylamide gel. For Cox4-mCherry, Dnm1-HA blots 8-10% gels were used. PVDF membranes were used for blotting, which were probed with anti-mCherry (1:5000), anti-HA (1:5000) antibodies, anti-Cox2 (1:1000). HRP conjugated secondary antibodies were used (1:5000) for western blotting and blots were developed using SuperSignal West PICO PLUS or FEMTO PLUS chemiluminescent substrate. Equal amount of proteins was loaded on a separate gel and stained with Coomassie Brillian Blue as a loading control.

### Immunocytochemistry

Mid-log phase cells were harvested and fixed in 1ml of 4% paraformaldehyde for 30 min at room temperature. After washing the cells twice with a phosphate-sorbitol buffer (0.1 M KPO4 and 1x in 0.1 M KPO4/1.2 M sorbitol), the pellet was resuspended in 1 ml of 0.1M KPO4/1.2M sorbitol. Cells were then spheroplasted with 10mg/ml Zymolyase for 10-30 min in presence of β-mercaptoethanol. Spheroplasting was checked by imaging cells on the microscope. For imaging, slides/dishes were prepared by washing in chilled methanol and 95% ethanol. Slides were then coated with 0.1% Poly-L lysine for 5-10 min and then washed thrice with dH2O. 15μl of spheroplast suspension was placed in each well for 5-10 min at room temperature and then aspirated, and immediately plunged into ice cold methanol for 5-6 min and then in cold acetone for 30 sec. Post air drying, 25μl of PBS-BSA was used for blocking. Primary antibody probing was done following 30 min of blocking, for 1 hr. After 5 washes with PBS-BSA, secondary antibody was added and incubated for 1-2 hr in a humid chamber in the dark. The wells were again washed with PBS-BSA five times (each wash for 10 min) and then twice with only PBS. Slides were mounted with mounted media (Slowfade DAPI reagent) and imaged.

## Acknowledgements

The authors are grateful to Dr. Ivan Ahel for sharing bacterial DarT and DarG constructs and Afroze Chimthanawala for assistance with SIM imaging. The authors thank Dr. Sunil Laxman and members of the AB and SL labs for helpful discussions and feedback on the manuscript. This work was supported by fellowships from TIFR (ND), CSIR (AS) as well as grants from HFSP CDA (00051/ 2017-C) (AB) and intramural funding from NCBS-TIFR (AB).

## Author contributions

ND: conception of project, experimental design, generation of tools and reagents, execution of experiments, data analysis, writing of manuscript. AS: experimental design, generation of tools and reagents, execution of experiments, data analysis, editing of manuscript. AB: conception of project, project supervision, experimental design and writing of manuscript.

## Supplementary information

### Supplementary figure legends

**Figure S1:**
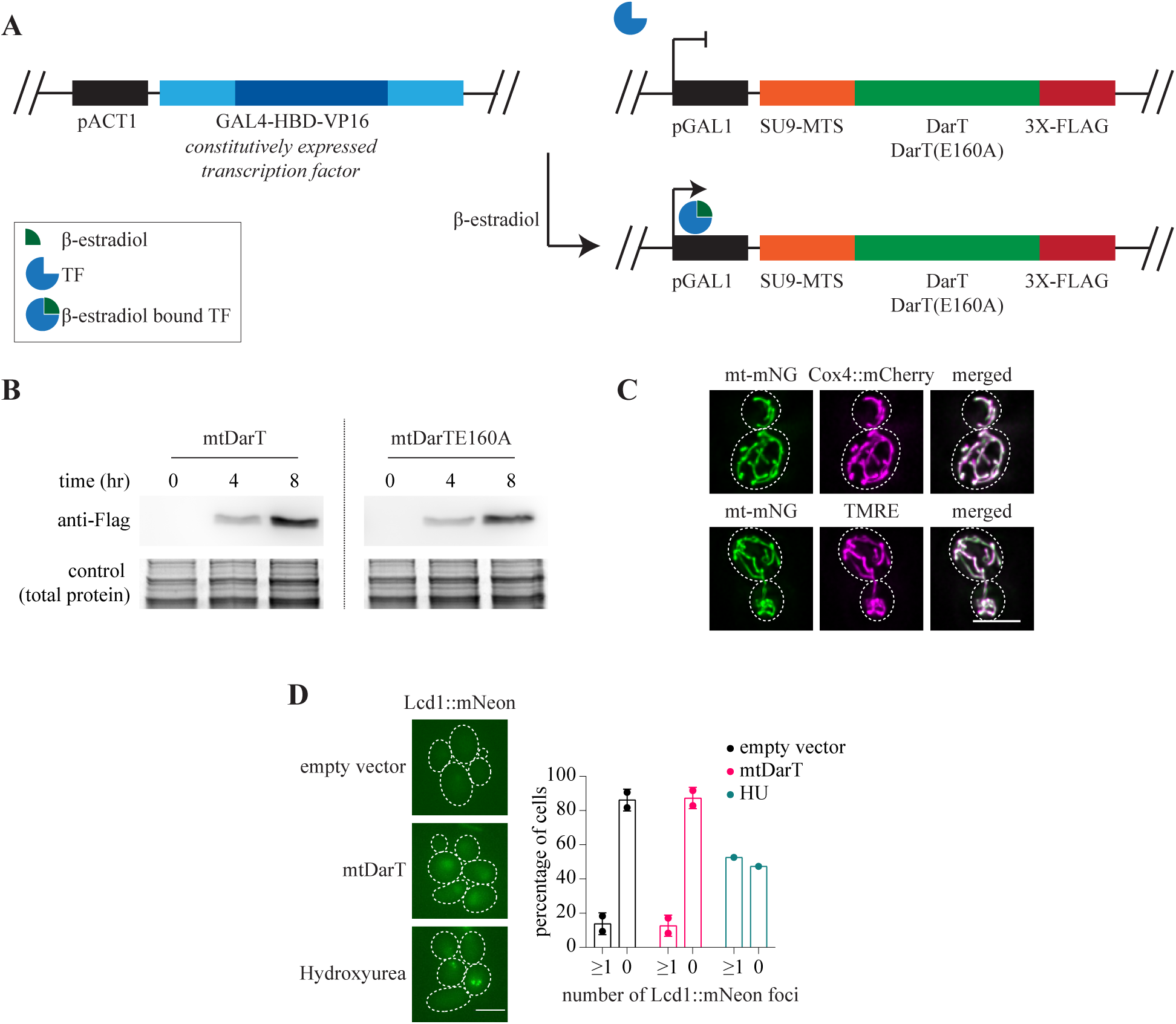
System to induce mitochondria-specific DNA damage. (A) Schematic of β-estradiol inducible system. Transcription factor GAL4-HBD-VP16 is constitutively expressed under a pACT1 promoter, and is activated on binding of β-estradiol. The activated transcription factor binds to pGAL1 and initiates transcription of gene under this promoter. (B) Western blots of FLAG-tagged mtDarT/TE160A after 4 and 8 hr of induction with 100nM β-estradiol. (C) Representative cells expressing mt-mNeongreen and Cox4::mCherry (top panel), and cell expressing mt-mNeongreen stained with TMRE (bottom panel). (D) Representative cells expressing Lcd1::mNeongreen post 6 hr of mtDarT induction or 2hr post Hydroxyurea (100mM) treatment. Percentage cells with or without Lcd1::mNeongreen foci is plotted. N>400 cells pooled from 2 biological replicates. (Scale bars-5μm, white dotted lines mark cell outlines, in all figures)

**Figure S2:**
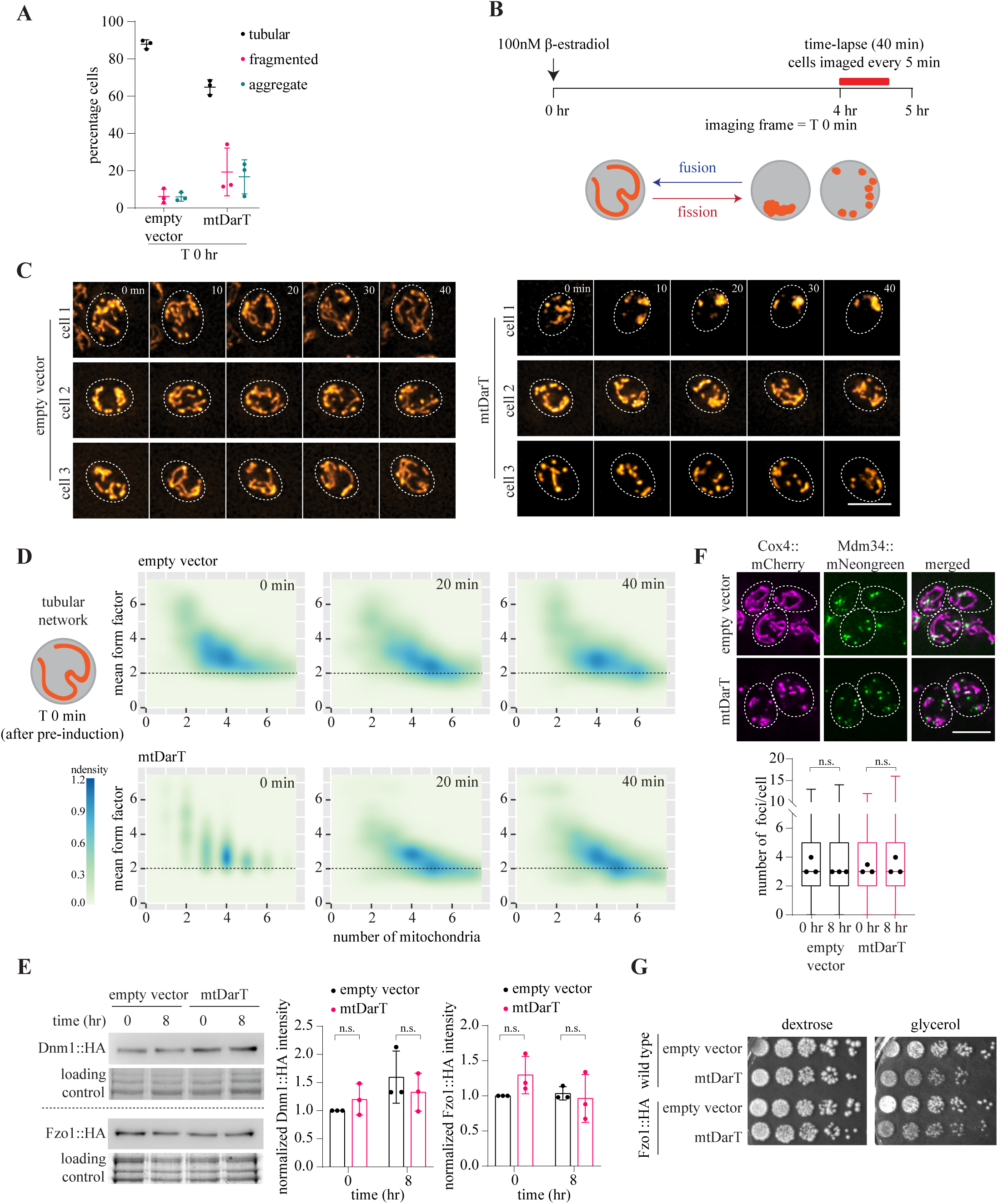
mtDNA damage impacts mitochondrial organization and dynamics. (A) Percentage of cells in each category in control (empty vector) or mtDarT cells at t=0hr (mean±SD from three biological replicates, n>150 in each replicate). (B) Schematic showing experimental design for time lapse experiments. (C) Montage of single cells expressing Cox4::mCherry tracked over time (mean form factor <2 at the beginning of the time lapse), imaged for 40 min after 4 hr of induction. T0 refers to the frame at the start of the imaging (after the period of pre-induction) here and in Fig. S2D. (D) Density plots with mean form factor and number of mitochondria over time, plotted for cells starting with MFF>2 at the beginning of time lapse; n=362 for empty vector, n=347 for mtDarT from 3 biological replicates. (E) Dnm1::HA and Fzo1::HA western blot with quantification from 3 biological replicates, normalized to empty vector at t=0; Error bars: SD. (F) Representative images of cells expressing Mdm34::mNeongreen and Cox4::mCherry after 8 hr of induction. Number of Mdm34 foci per cell are plotted from 3 biological replicates; n>680. (G) Cell survival on plates containing either dextrose or glycerol as a carbon source, to ensure C-terminal tagging of Fzo1 does not affect cell growth. Statistical significance in E was calculated using unpaired Student’s t-test. (*** p<0.0001, ** p<0.005, * p< 0.05, n.s.- not significant, in all cases).

**Figure S3:**
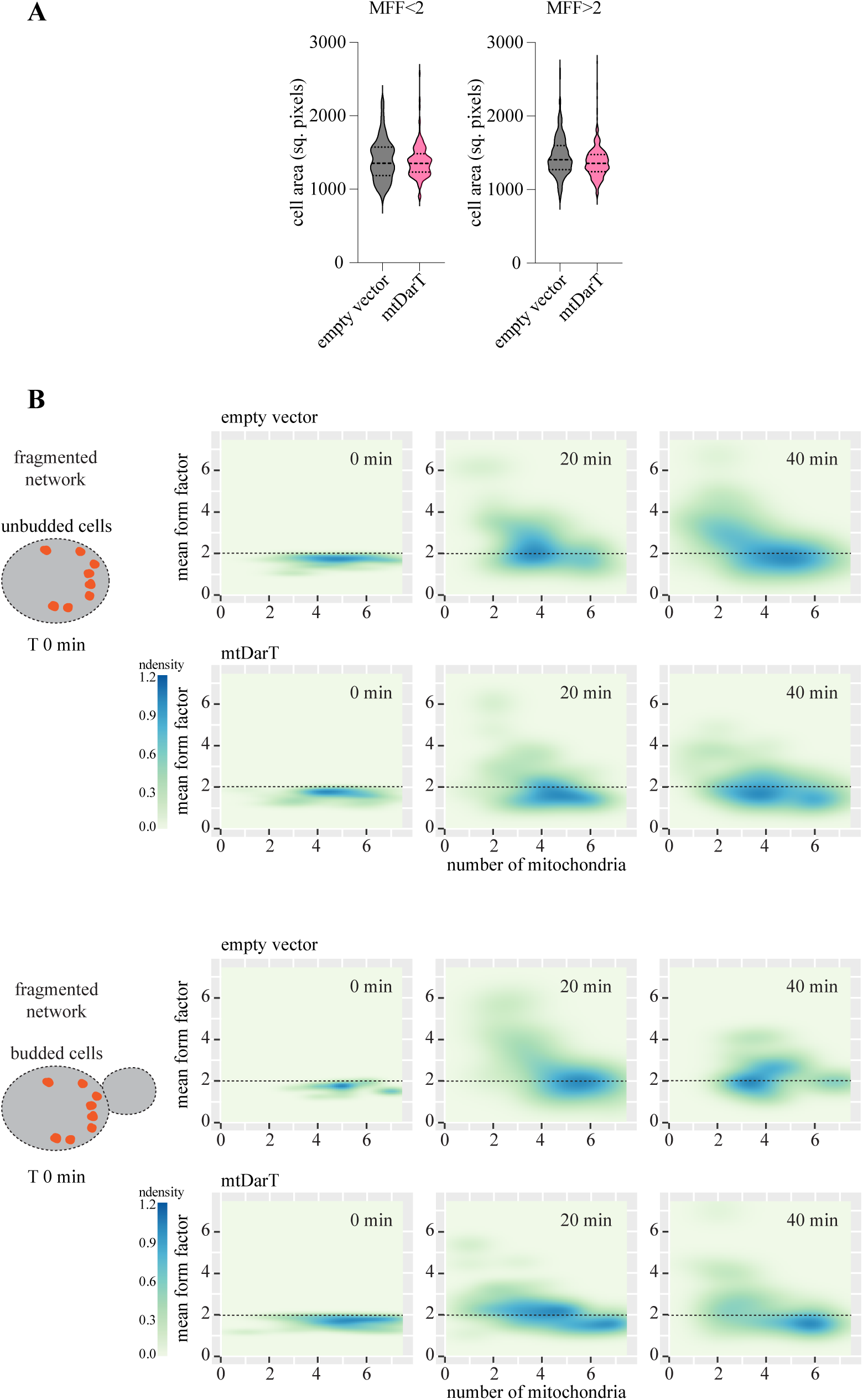
mtDNA damage impacts mitochondrial organization and dynamics. (A) Cell area plotted as violin plots for all the cells in Fig 2 and S2D at t=0 min. (B) Density plots with Mean form factor and number of mitochondria over time, plotted for cells in Fig 2F, classified based on presence or absence of buds. T0 refers to the frame at the start of the imaging (after the period of pre-induction)

**Figure S4:**
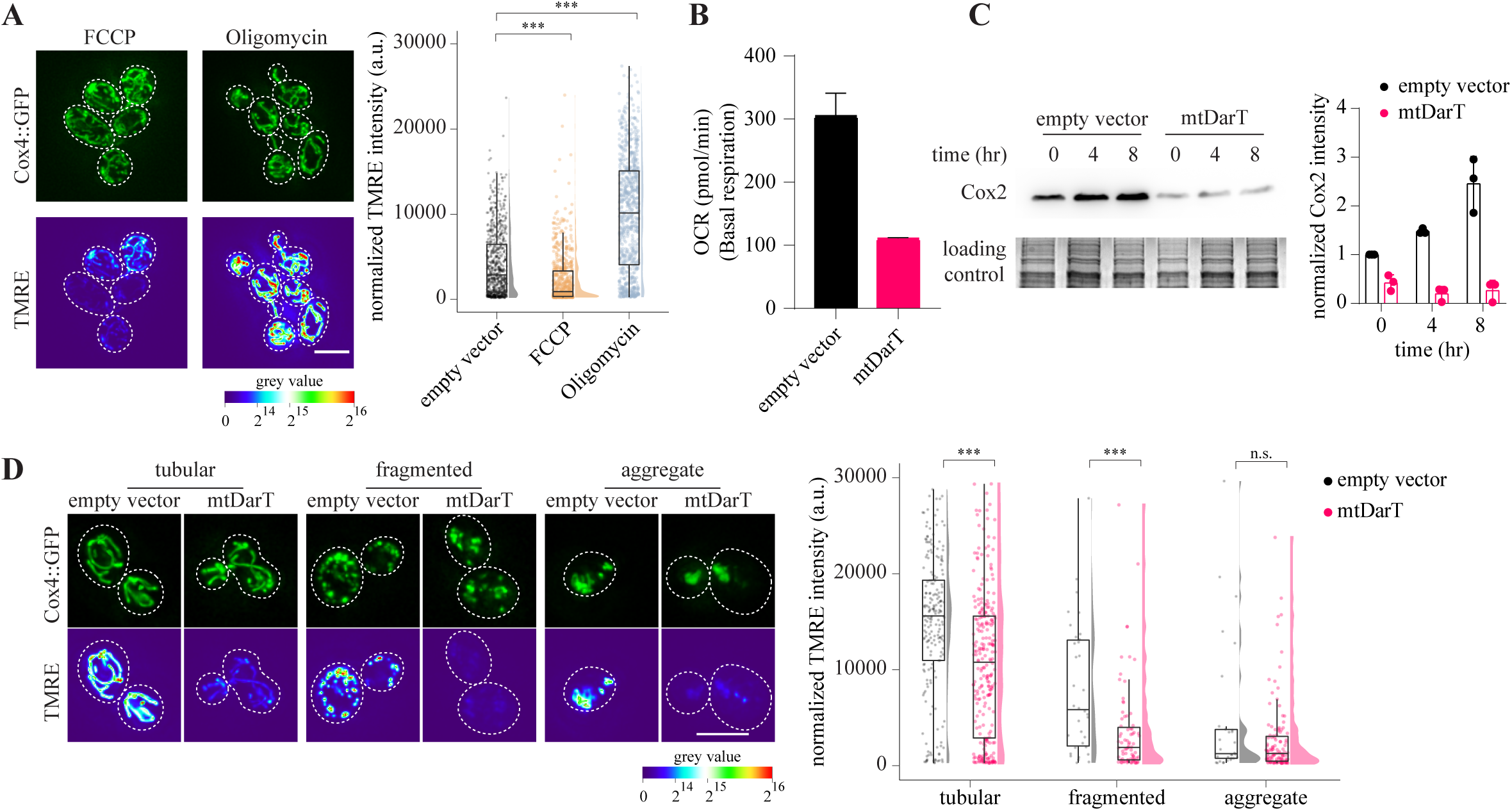
mtDNA damage results in mitochondrial dysfunction. (A) Representative images of cells expressing Cox4::GFP stained with TMRE after treatment with FCCP (2 μM) for 20 min or Oligomycin (0.5 μg/ml) for 4 hr (left panel) (Cox4::GFP: green). Total TMRE intensity normalized to cell area is plotted; n>650 from 3 biological replicates for FCCP and Oligomycin and 2 biological replicates for empty vector (right panel). (B) Oxygen consumption rate (OCR) estimated using Seahorse assay post 4hr of induction. Mean and SD plotted from technical replicates. (C) Cox2 western blot with quantification from 3 biological replicates, normalized to empty vector at t=0; Error bars: SD. (D) Representative images of cells expressing COX4::GFP stained with TMRE after 4 hr of induction with 100nM β-estradiol. Examples show cells from three different morphological categories for both empty vector and mtDarT (top panel). Total TMRE intensity normalized to cell area is plotted for cells with three mitochondrial morphologies, after 4 hr of induction (for empty vector, n=215 for tubular, n=35 for fragmented, n=30 aggregate; for mtDarT, n=258 for tubular, n=79 for fragmented, n=118 for aggregate). Statistical significance in A, D was calculated using Mann-Whitney U test.

**Figure S5:**
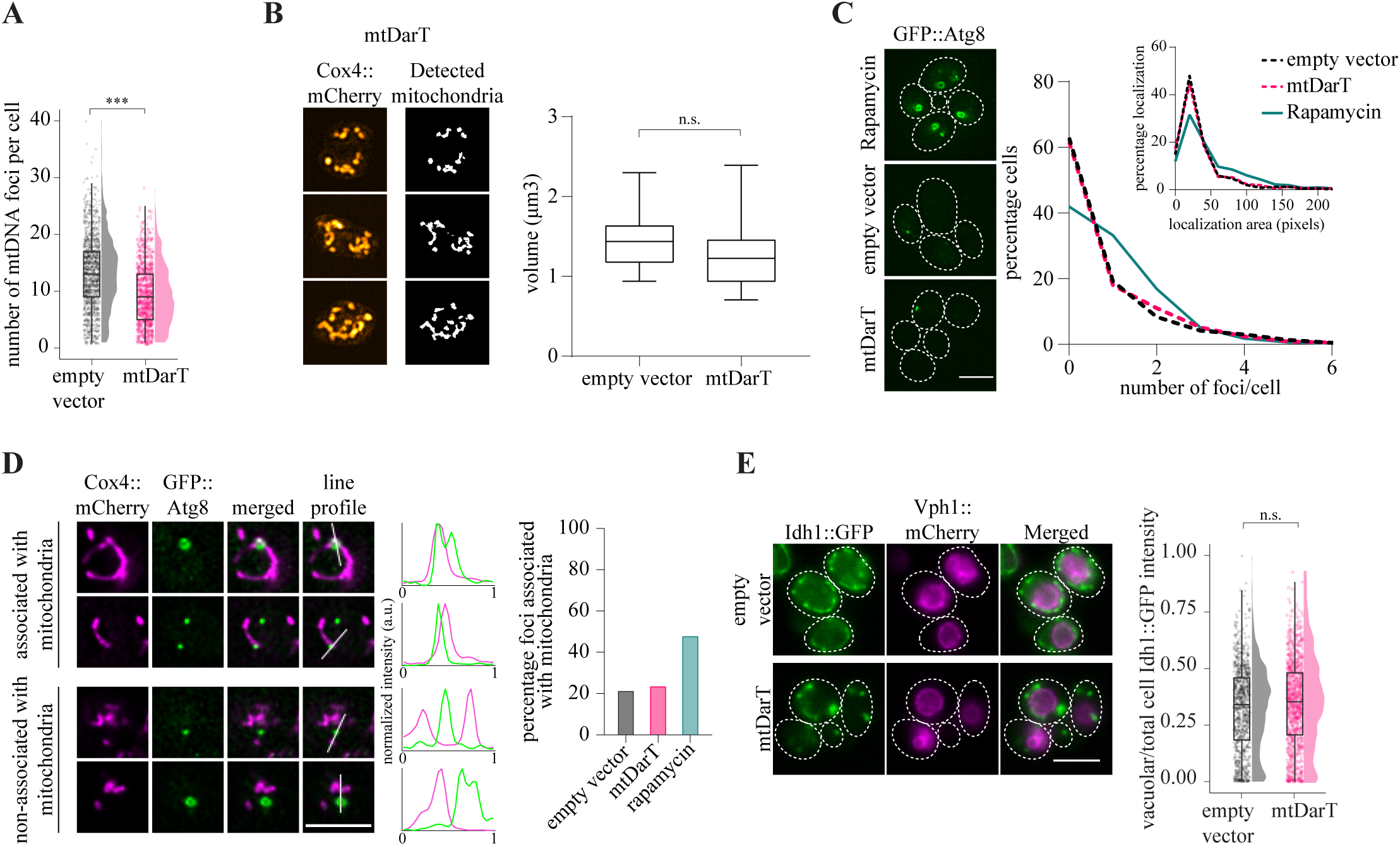
Changes in mitochondrial morphology accompany loss of mtDNA but not mitochondria. (A) Number of mtDNA foci per cell after 8 hr of induction is plotted; n>930 from 3 biological replicates. (B) Representative cells imaged with Structured Illumination Microscopy, along with segmentation output after Mitograph analysis (left panel). Mitochondrial volume was analysed via estimation of number of voxels detected using Mitograph, each data point represents one cell; n=12 for empty vector and n=24 for mtDarT. (C) Representative images of cells expressing GFP::Atg8 under the endogenous *ATG8* promoter treated with Rapamycin (20nM) for 4 hr and cells transformed with empty vector or mtDarT after 8 hs of induction with 100nM β-estradiol (left panel). Frequency distribution of number of GFP::Atg8 localization (right panel). (D) Representative cells showing GFP::Atg8 localizations associated or non-associated with mitochondria. Percentage localizations associated with mitochondria is plotted post 8 hr of induction, or 4 hr of rapamycin (20nM) treatment. Data is pooled from 2 biological replicates. N>90 localizations. (E) Vacuolar intensity of Idh1::GFP normalized to total cell intensity after 4 hr of induction with 100nM β-estradiol; n>830 from 2 biological replicates (left panel), with representative images of cells expressing Idh1::GFP and Vph1::mCherry (right panel). Statistical significance in A,B,E was calculated using Mann-Whitney U test.

**Figure S6:**
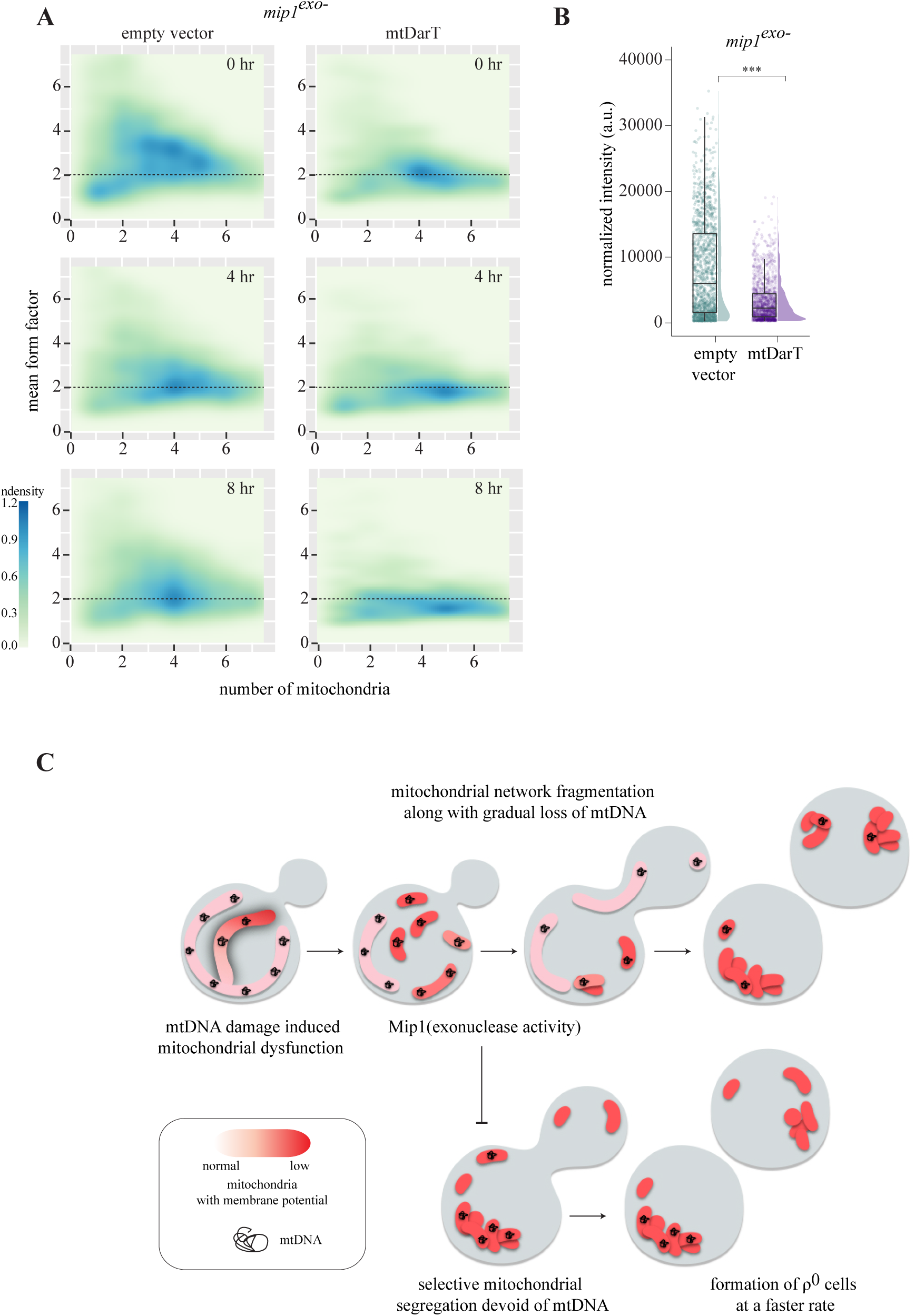
Exonuclease activity of the mtDNA polymerase, Mip1, is essential for mtDNA maintenance. (A) Density plots with mean form factor and number of mitochondria over time for *mip1^exo-^* cells; n>710 in each plot, from 3 biological replicates. (B) Total TMRE intensity normalized to cell area for *mip1^exo-^* cells is plotted; n>1350 from 3 biological replicates. (C) Schematic summarizing key observations of this study. mtDNA damage triggers mitochondrial dysfunction and associated changes in mitochondrial organization. Exonuclease function of replicative polymerase, Mip1, is required to prevent rapid loss of mtDNA under damage. For details, see discussion. Statistical significance in B was calculated using Mann-Whitney U test.

**Supplementary Table S1:**
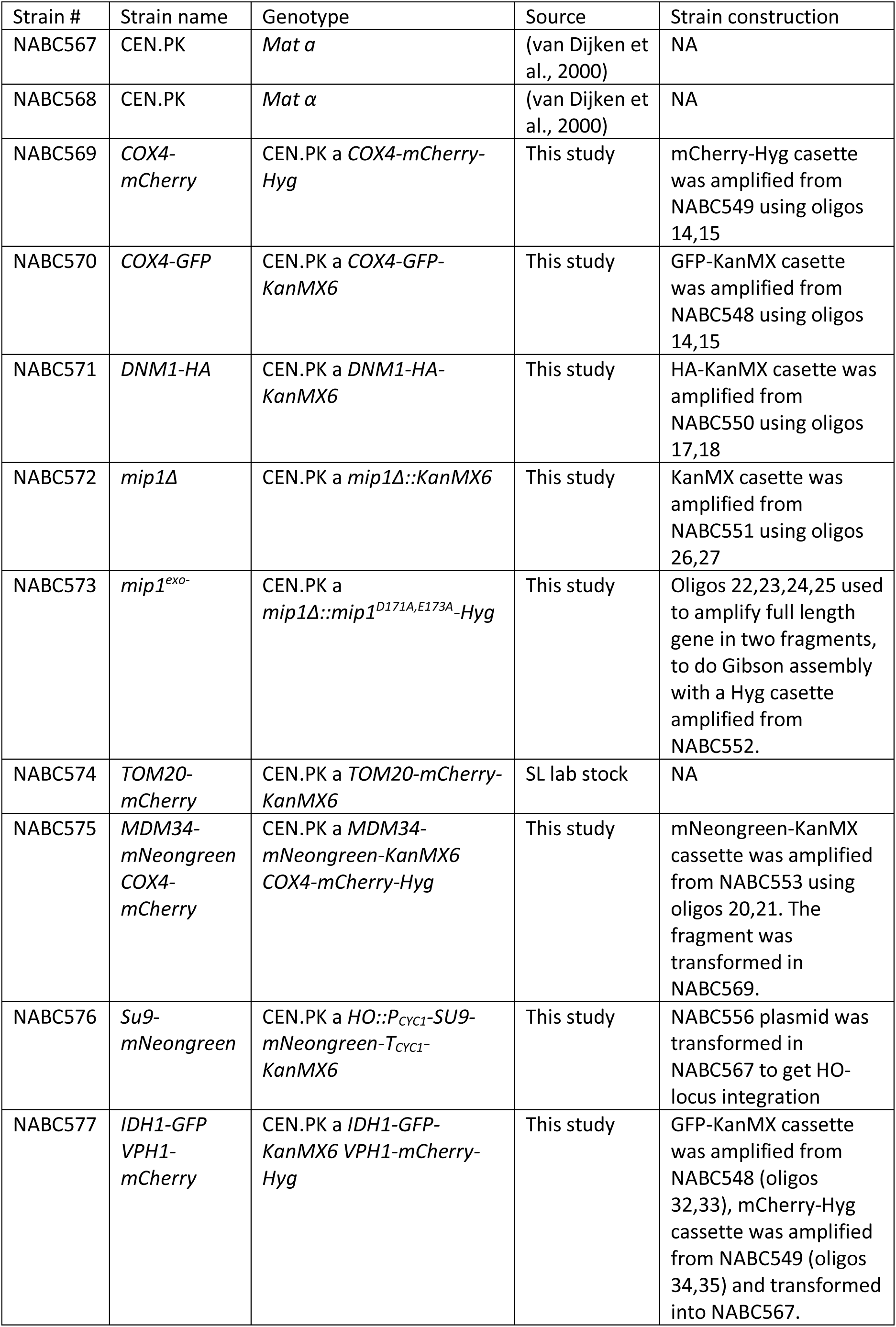
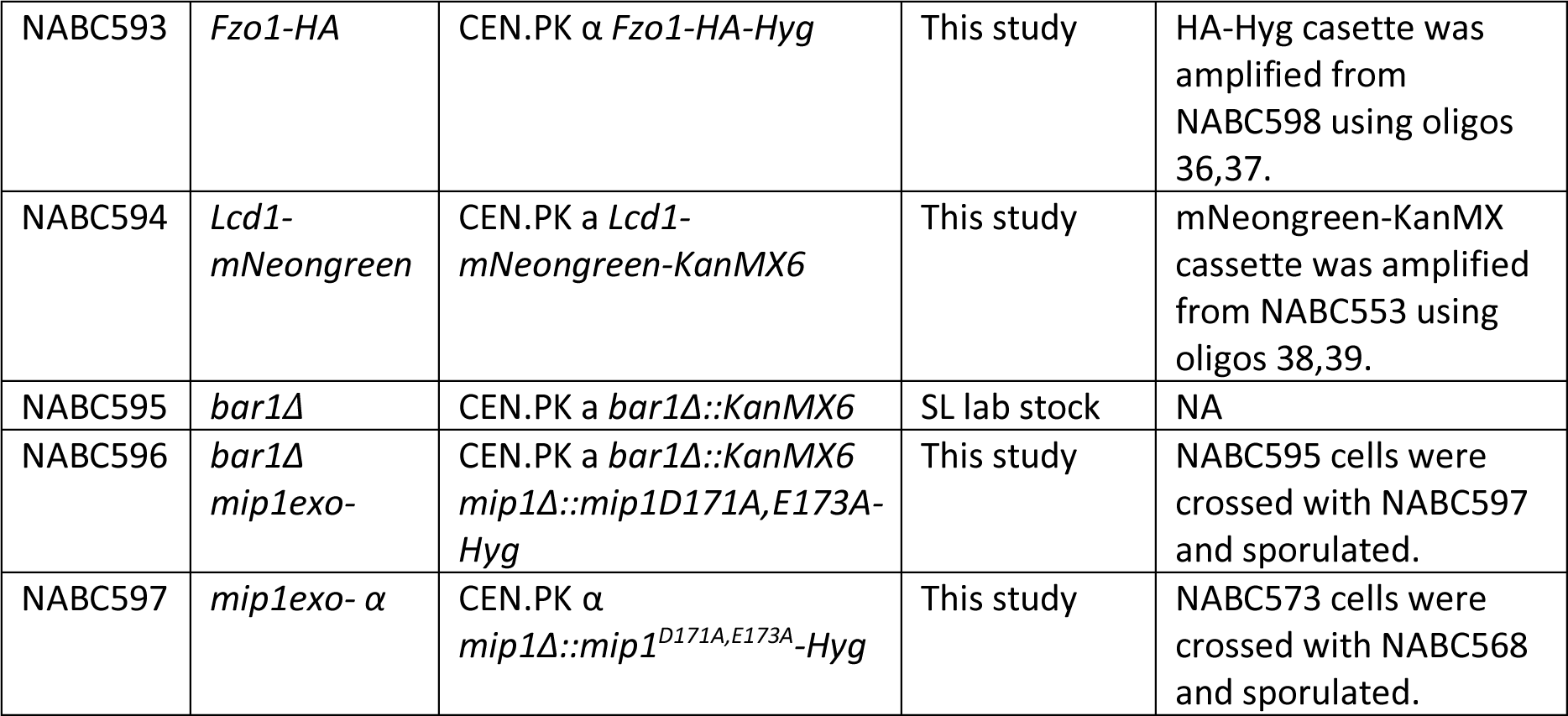
Strains used in present study.

**Supplementary Table S2:**
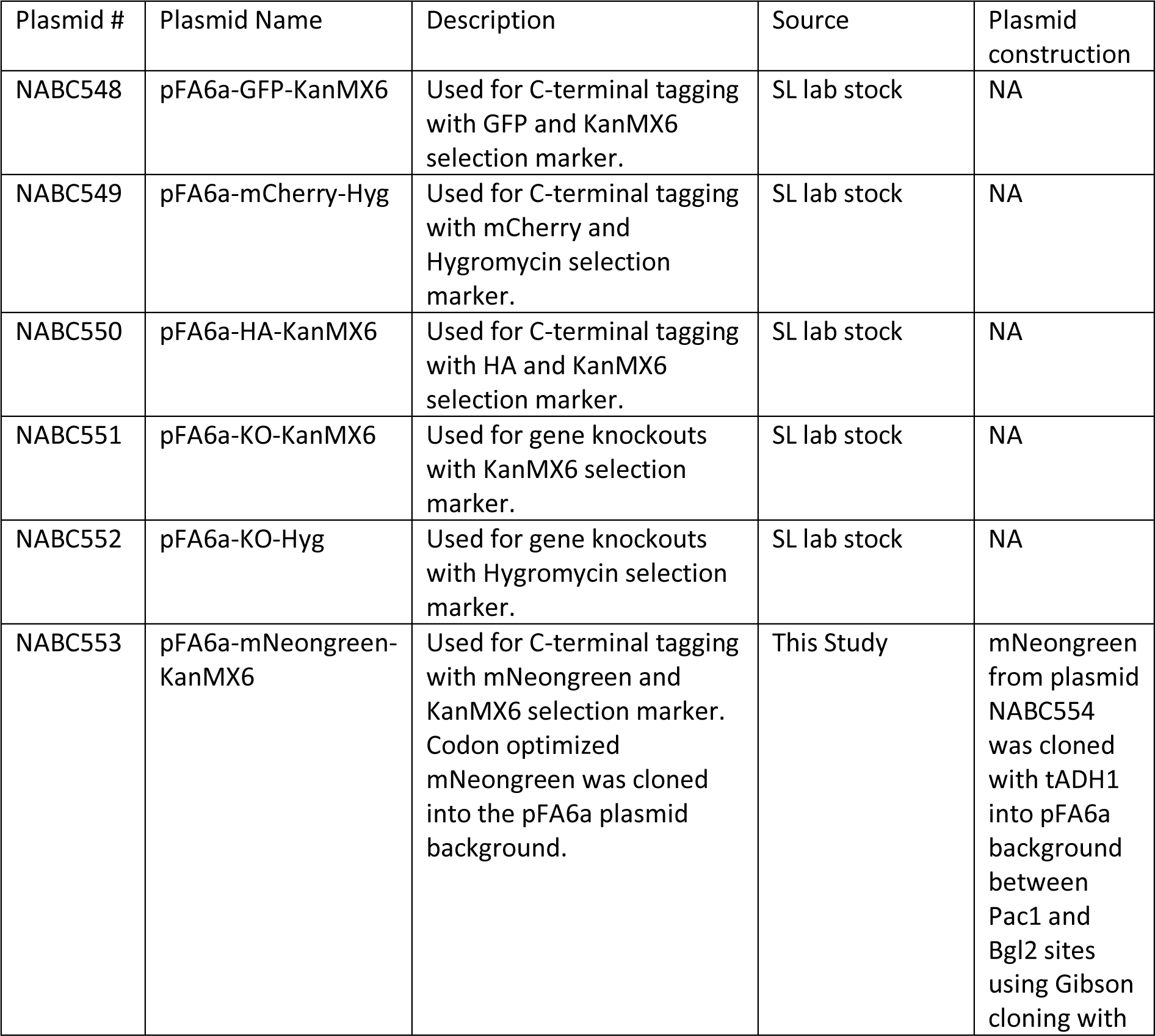
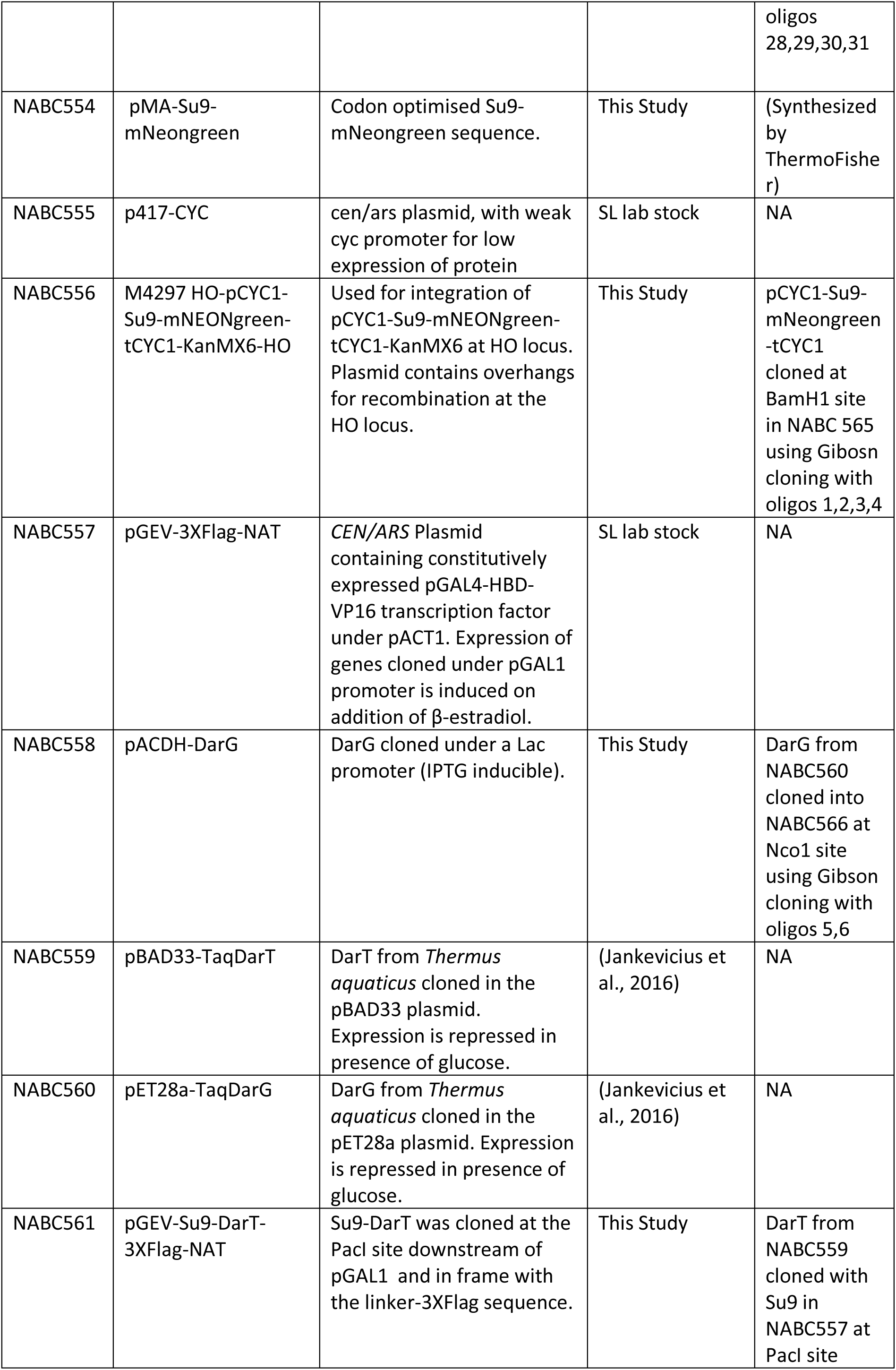
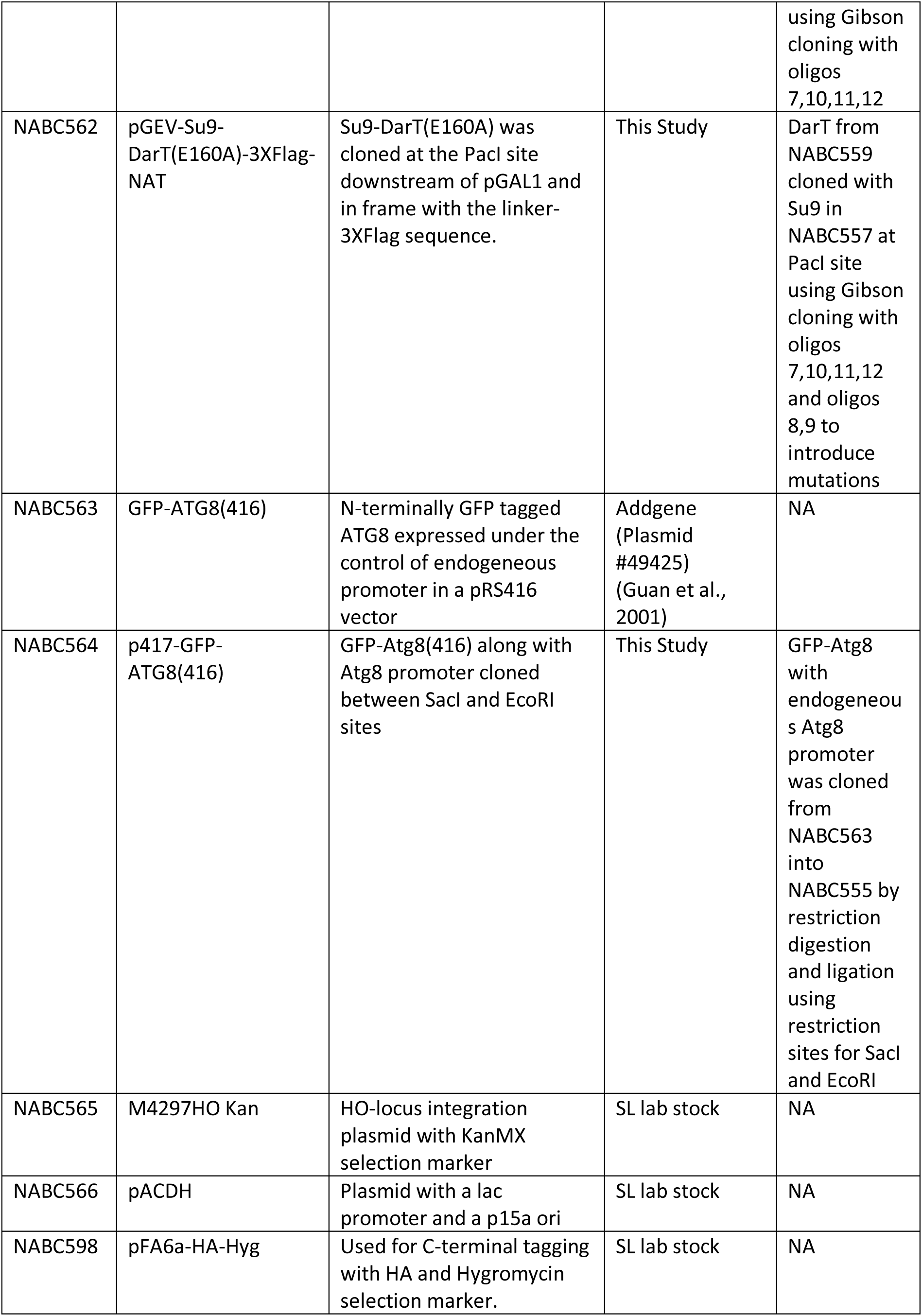
Plasmids used in present study.

| Plasmid # | Plasmid Name | Description | Source | Plasmid construction |
| --- | --- | --- | --- | --- |
| NABC548 | pFA6a-GFP-KanMX6 | Used for C-terminal tagging with GFP and KanMX6 selection marker. | SL lab stock | NA |
| NABC549 | pFA6a-mCherry-Hyg | Used for C-terminal tagging with mCherry and Hygromycin selection marker. | SL lab stock | NA |
| NABC550 | pFA6a-HA-KanMX6 | Used for C-terminal tagging with HA and KanMX6 selection marker. | SL lab stock | NA |
| NABC551 | pFA6a-KO-KanMX6 | Used for gene knockouts with KanMX6 selection marker. | SL lab stock | NA |
| NABC552 | pFA6a-KO-Hyg | Used for gene knockouts with Hygromycin selection marker. | SL lab stock | NA |
| NABC553 | pFA6a-mNeongreen-KanMX6 | Used for C-terminal tagging with mNeongreen and KanMX6 selection marker. Codon optimized mNeongreen was cloned into the pFA6a plasmid background. | This Study | mNeongreen from plasmid NABC554 was cloned with tADH1 into pFA6a background between Pac1 and Bgl2 sites using Gibson cloning with oligos<br>28,29,30,31 |
| NABC554 | pMA-Su9-mNeongreen | Codon optimised Su9-mNeongreen sequence. | This Study | (Synthesized by ThermoFisher) |
| NABC555 | p417-CYC | cen/ars plasmid, with weak cyc promoter for low expression of protein | SL lab stock | NA |
| NABC556 | M4297 HO-pCYC1-Su9-mNEONGreen-tCYC1-KanMX6-HO | Used for integration of pCYC1-Su9-mNEONGreen-tCYC1-KanMX6 at HO locus. Plasmid contains overhangs for recombination at the HO locus. | This Study | pCYC1-Su9-mNeongreen-tCYC1 cloned at BamH1 site in NABC 565 using Gibson cloning with oligos 1,2,3,4 |
| NABC557 | pGEV-3XFlag-NAT | CEN/ARS Plasmid containing constitutively expressed pGAL4-HBD-VP16 transcription factor under pACT1. Expression of genes cloned under pGAL1 promoter is induced on addition of β-estradiol. | SL lab stock | NA |
| NABC558 | pACDH-DarG | DarG cloned under a Lac promoter (IPTG inducible). | This Study | DarG from NABC560 cloned into NABC566 at Nco1 site using Gibson cloning with oligos 5,6 |
| NABC559 | pBAD33-TaqDarT | DarT from *Thermus aquaticus* cloned in the pBAD33 plasmid. Expression is repressed in presence of glucose. | (Jankevicius et al., 2016) | NA |
| NABC560 | pET28a-TaqDarG | DarG from *Thermus aquaticus* cloned in the pET28a plasmid. Expression is repressed in presence of glucose. | (Jankevicius et al., 2016) | NA |
| NABC561 | pGEV-Su9-DarT-3XFlag-NAT | Su9-DarT was cloned at the PacI site downstream of pGAL1 and in frame with the linker-3XFlag sequence. | This Study | DarT from NABC559 cloned with Su9 in NABC557 at PacI site using Gibson cloning with oligos 7,10,11,12 |
| NABC562 | pGEV-Su9-DarT(E160A)-3XFlag-NAT | Su9-DarT(E160A) was cloned at the PacI site downstream of pGAL1 and in frame with the linker-3XFlag sequence. | This Study | DarT from NABC559 cloned with Su9 in NABC557 at PacI site using Gibson cloning with oligos 7,10,11,12 and oligos 8,9 to introduce mutations |
| NABC563 | GFP-ATG8(416) | N-terminally GFP tagged ATG8 expressed under the control of endogeneous promoter in a pRS416 vector | Addgene (Plasmid #49425) (Guan et al., 2001) | NA |
| NABC564 | p417-GFP-ATG8(416) | GFP-Atg8(416) along with Atg8 promoter cloned between SacI and EcoRI sites | This Study | GFP-Atg8 with endogeneous Atg8 promoter was cloned from NABC563 into NABC555 by restriction digestion and ligation using restriction sites for SacI and EcoRI |
| NABC565 | M4297HO Kan | HO-locus integration plasmid with KanMX selection marker | SL lab stock | NA |
| NABC566 | pACDH | Plasmid with a lac promoter and a p15a ori | SL lab stock | NA |
| NABC598 | pFA6a-HA-Hyg | Used for C-terminal tagging with HA and Hygromycin selection marker. | SL lab stock | NA |

**Supplementary Table S3:**
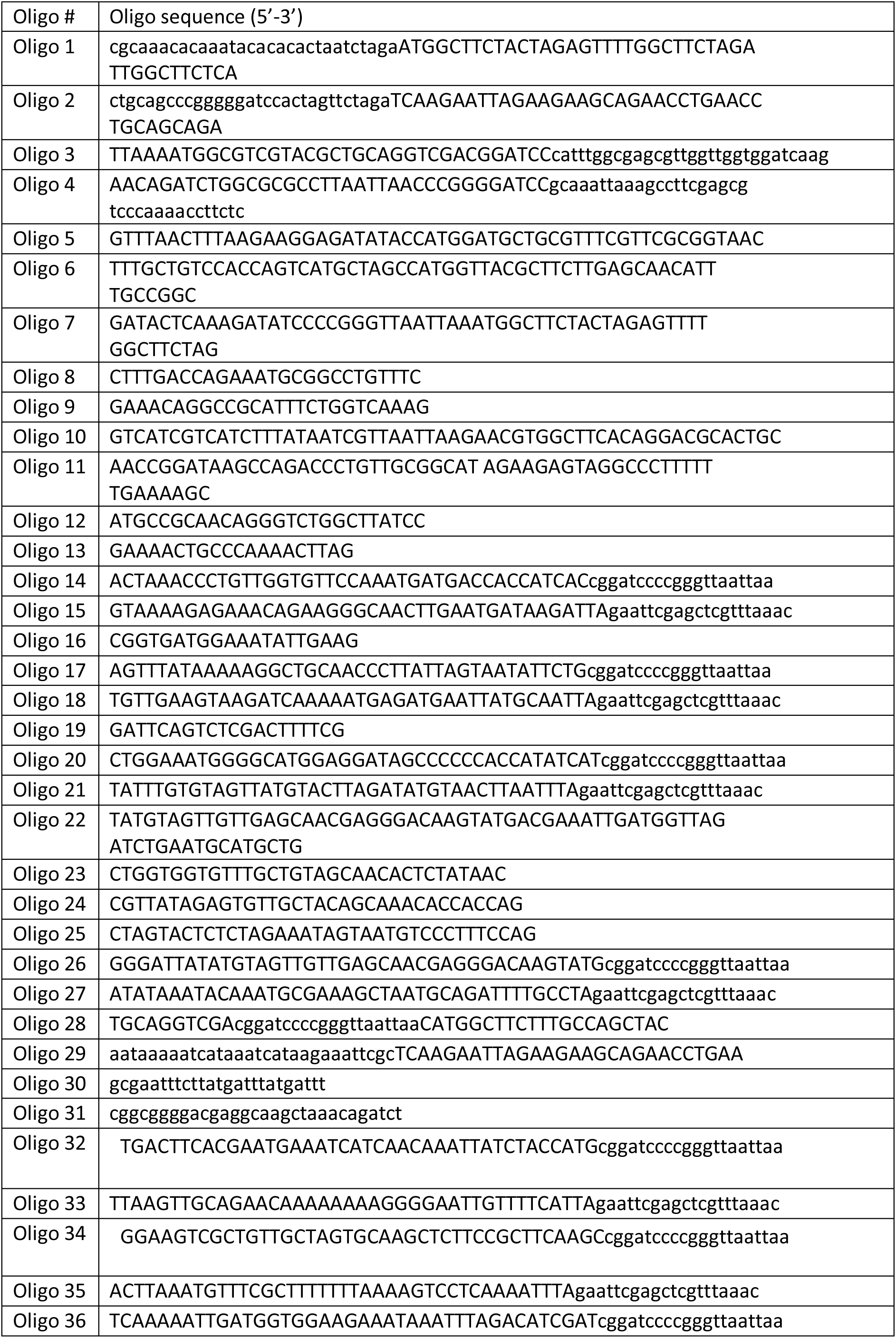
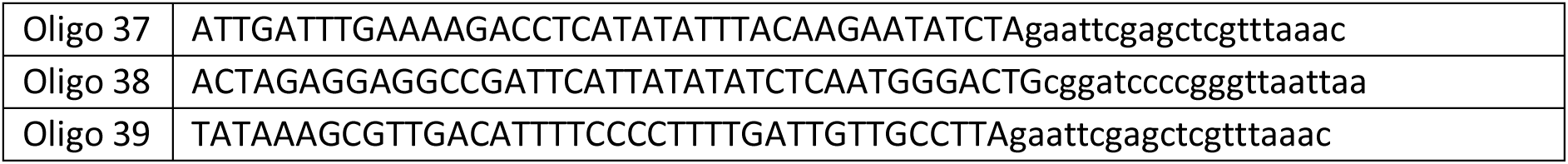
Oligos used in present study.

